# Pyruvate kinase modulates the link between β-cell fructose metabolism and insulin secretion

**DOI:** 10.1101/2024.08.15.608033

**Authors:** Naoya Murao, Risa Morikawa, Yusuke Seino, Kenju Shimomura, Yuko Maejima, Tamio Ohno, Norihide Yokoi, Yuichiro Yamada, Atsushi Suzuki

## Abstract

2

Glucose triggers insulin secretion from pancreatic β-cells through intracellular glucose metabolism, ATP production, and closure of ATP-sensitive K^+^ channels (K_ATP_ channels). Fructose also stimulates insulin secretion, but the underlying mechanisms remain unclear. This study investigated the contribution of phospholipase C (PLC) signaling and fructose metabolism to fructose-stimulated insulin secretion (FSIS) using MIN6-K8 clonal β-cells and mouse islets.

Fructose-induced PLC activation, assessed by inositol 1-phosphate accumulation, was reduced in fructose-unresponsive β-cell models, such as diabetic mouse islets and K_ATP_ channel-deficient β-cells, suggesting that β-cell fructose responsiveness is primarily determined by PLC signaling.

Although FSIS was dependent on K_ATP_ channels and Ca^2+^ influx, the ATP/ADP ratio was unexpectedly lowered by fructose, and suppression of intracellular fructose metabolism hardly affected FSIS. Metabolic flux analysis revealed that the accumulation of fructose 1-phosphate (F1P) suppressed pyruvate kinase (PK) activity, contributing to ATP depletion. Strikingly, a small-molecule PK activator, TEPP-46, antagonized F1P-mediated PK suppression, prevented the drop in the ATP/ADP ratio, and restored FSIS in MIN6-K8 cells, normal mouse islets, and fructose-unresponsive diabetic mouse islets.

These findings revealed the metabolic effects of fructose in β-cells and identified PK as a key regulator linking β-cell fructose metabolism and FSIS, thereby providing new insights into the mechanisms of insulin secretion and potential therapeutic targets for fructose-associated metabolic diseases.

**GRAPHICAL ABSTRACT:** 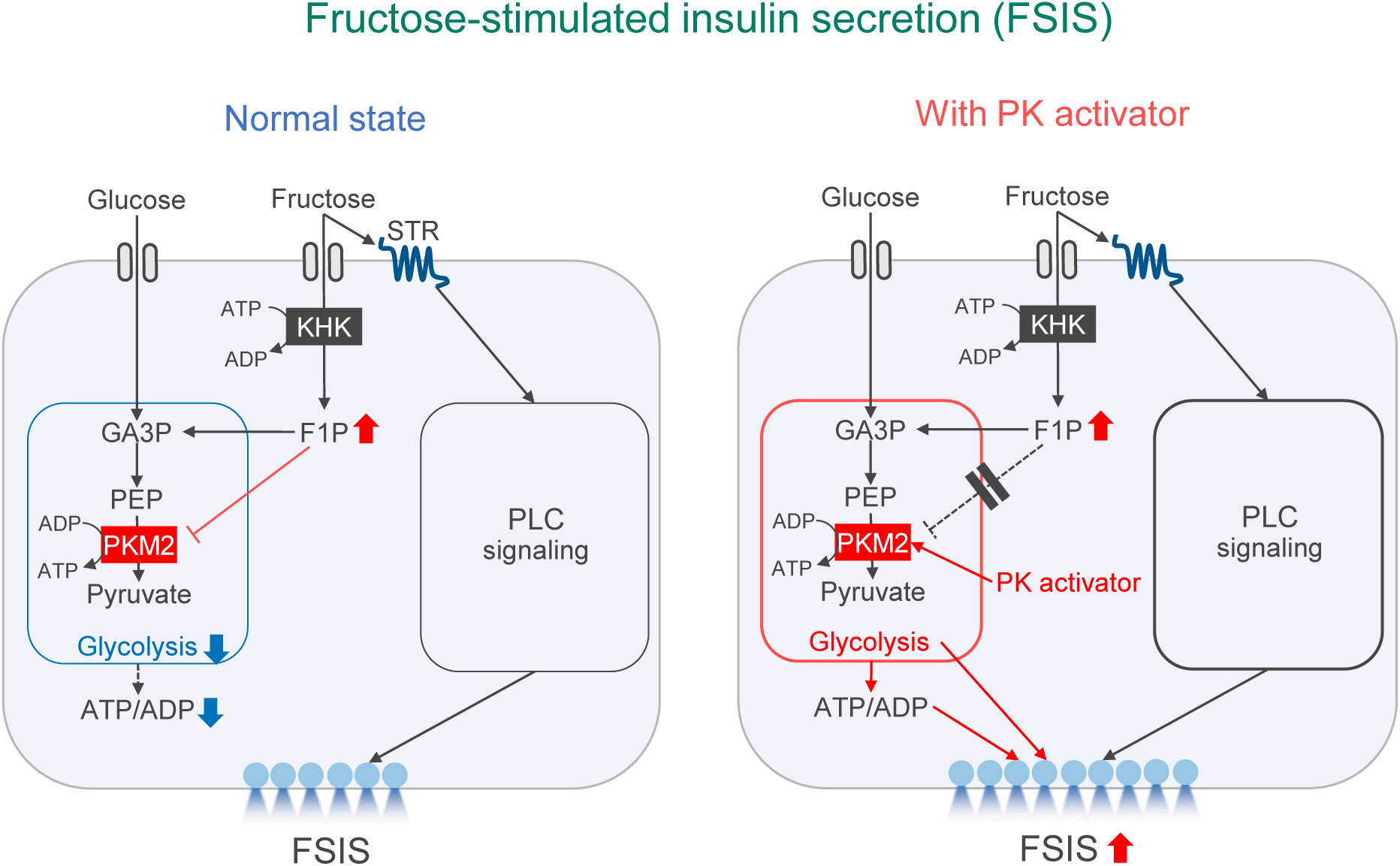

Left: Fructose-stimulated insulin secretion (FSIS) is driven by sweet taste receptor (STR)-mediated PLC signaling in pancreatic β-cells. Meanwhile, fructose metabolism does not promote FSIS because fructose causes accumulation of fructose 1-phosphate (F1P), which suppresses pyruvate kinase M2 (PKM2), lowering the ATP/ADP ratio.

Right: A small-molecule PK activator counteracted F1P-mediated PKM2 inhibition, prevented ATP decrease, and substantially enhanced FSIS in normal and diabetic mouse β-cells. Thus, PK has been identified as a key regulator linking β-cell fructose metabolism and FSIS.

## 5 INTRODUCTION

Dietary monosaccharides, such as glucose, fructose, and mannose, stimulate pancreatic β-cells to increase insulin secretion (Grodsky et al., 1963; Ashcroft et al., 1972). The process of glucose-stimulated insulin secretion (GSIS) is critical for glucose homeostasis, and numerous studies have delineated its molecular mechanisms. It is well established that GSIS is regulated by β-cell glucose metabolism through two signaling pathways: the triggering and amplifying pathways (Henquin, 2000; Henquin, 2009). The triggering pathway refers to the increase in intracellular [Ca^2+^] after a sequence of events: an increase in intracellular ATP levels, closure of ATP-sensitive K^+^ (K_ATP_) channels, membrane depolarization, and opening of voltage-dependent Ca^2+^ channels (VDCCs). The amplifying pathway augments the triggering pathway only when the latter is already active through various glucose-related metabolites such as adenine nucleotides, citrate, GTP, and glutamate (Prentki et al., 2013; Murao et al., 2017; Han et al., 2021).

Fructose is the second most abundant monosaccharide in the diet after glucose (Larke et al., 2023). It is widely found in fruits and certain sweeteners, making it a significant source of energy for many people. Fructose has been known to stimulates insulin secretion at physiological glucose levels (Grodsky et al., 1963; Ashcroft et al., 1972; Zawalich et al., 1977; Grant et al., 1980). However, the mechanism of fructose-stimulated insulin secretion (FSIS) is less well understood than that of glucose-stimulated insulin secretion (GSIS).

In various cell types, fructose is transported primarily through GLUT5 (SLC2A5) and is converted to F1P by ketohexokinase (KHK), the first rate-limiting enzyme in fructolysis (Hannou et al., 2018). It is then metabolized to glyceraldehyde 3-phosphate (GA3P) and dihydroxyacetone phosphate (DHAP) and enters glycolytic/gluconeogenic metabolite pool, part of which is used for ATP production (Hannou, 2018, see also Figure 2).

Thus, it is reasonable to assume that ATP production by fructose metabolism stimulates FSIS in a pathway similar to that of glucose. However, this relationship has not yet been thoroughly investigated. To address this, we examined β-cell fructose metabolism and its functional relationship with insulin secretion using clonal β-cell lines and mouse islets as a model. Unexpectedly, we found that fructose metabolism in β-cells did not contribute to FSIS. Through detailed metabolic profiling, we discovered that fructose inhibits pyruvate kinase (PK) and hinders glycolysis. Remarkably, we showed that PK activators counteracted this inhibition, enhancing FSIS, which was also effective in fructose-unresponsive diabetic mouse islets.

## 6 MATERIALS AND METHODS

### 6.1. Animals

C57BL/6JJcl (RRID:IMSR_JCL:JCL:MIN-0003) mice were purchased from CLEA Japan (Tokyo, Japan) and used for experiments at 19–20 weeks of age. NSY.B6-*Tyr*+, *A^y^*/Hos mice were a gift from Hoshino Laboratory Animals (Ibaraki, Japan) and were used for experiments at 19–20 weeks of age. Mice were maintained under specific-pathogen-free conditions at 23 ± 2 °C and 55 ± 10% relative humidity with 12-hour light-dark cycles (8am–8pm), with free access to water and standard chow CE-2 (CLEA Japan, Tokyo, Japan). The health status of the mice was checked regularly. All experiments were performed using male mice. Body weight and blood glucose levels were measured at 9am in the ad-lib-fed state. The number of mice analyzed is indicated in the figure legends. All animal experiments were performed with the approval APU22080 by the Institutional Animal Care and Use Committee of Fujita Health University, complying with the Guidelines for Animal Experimentation at Fujita Health University and current Japanese guidelines and regulations for scientific and ethical animal experimentation (Kagiyama et al., 2006).

### 6.2. Cell lines

MIN6-K8 cells were generated as previously described (Iwasaki et al., 2010) and were kindly provided by Professor Junichi Miyazaki (Osaka University). *Kcnj11^-/-^* β-cells (clone *Kcnj11*^-/-^ βCL1) are β-cells deficient in Kir6.2, a pore-forming subunit of the K_ATP_ channel. They were established by sub-cloning MIN6-K8 cells transfected with Cas9 nickase and guide RNA pairs targeting mouse *Kcnj11,* as described previously (Oduori et al., 2020). All cells were cultured in Dulbecco’s modified Eagle’s medium (DMEM) containing 4500 mg/L glucose (Sigma-Aldrich, St. Louis, MO, USA, Cat# D5796) supplemented with 10% fetal bovine serum (FBS) (BioWest, Nuaillé, France, Cat# S1400-500) and 5 ppm 2-mercaptoethanol. The cells were maintained at 37 °C with 5% CO_2_.

### 6.3. Plasmids

GW1-PercevalHR was a gift from Gary Yellen (Addgene plasmid # 49082; http://n2t.net/addgene:49082; RRID: Addgene_49082) (Tantama et al., 2013).

### 6.4. Reagents

Krebs-Ringer bicarbonate buffer-HEPES (133.4 mM NaCl, 4.7 mM KCl, 1.2 mM KH_2_PO_4_, 1.2 mM MgSO_4_, 2.5 mM CaCl_2_, 5 mM NaHCO_3_, 10 mM HEPES) supplemented with 0.1% bovine-serum albumin (Sigma-Aldrich, St. Louis, MO, USA, Cat# A6003) and 2.8 mM glucose (2.8G-KRBH) adjusted to pH 7.4 was used in in vitro experiments. Additional glucose (final concentration: 8.8 mM), U73122 (CAS: 112648-68-7, Cat# 70740), and fructose were added to KRBH during the stimulation period. Nifedipine (FUJIFILM Wako Pure Chemical, Osaka, Japan, Cat# 14505781) and TEPP-46 (Selleck Chemicals, Houston, TX, USA, CAS: 1221186-53-3, Cat# S7302) were added during the pre-incubation and stimulation periods. The reagents used for stimulation were stored as a 1000× concentrate in dimethyl sulfoxide (DMSO) (FUJIFILM Wako Pure Chemical, Osaka, Japan, Cat# 041-29351) and diluted with KRBH shortly before the experiment. An equal volume of DMSO was added to the vehicle control.

### 6.5. Isolation of pancreatic islets from mice

Digesting solution was formulated by supplementing 0.1w/v% Collagenase from Clostridium histolyticum (Sigma-Aldrich, St. Louis, MO, USA, Cat# C6885) to Hanks’ balanced salt solution (136.9 mM NaCl, 5.4 mM KCl, 0.8 mM MgSO_4_, 0.3 mM Na_2_HPO_4_, 0.4 mM KH_2_PO_4_, 4.2 mM NaHCO_3_, 10 mM HEPES, 1.3 mM CaCl_2_, 2 mM glucose). The mice were euthanized by isoflurane exposure. Pancreas was digested by 10-min incubation at 37 °C following intraductal injection of digesting solution. Islets were hand-picked and separated from exocrine tissues, transferred to 60-mm non-treated plates (AGC Techno Glass, Shizuoka, Japan, Cat# 1010-060), and cultured overnight in RPMI-1640 (Sigma-Aldrich, St. Louis, MO, USA, Cat# R8758) supplemented with 10% FBS (BioWest, Nuaillé, France, Cat# S1400-500) and 1% penicillin-streptomycin solution (FUJIFILM Wako Pure Chemical, Osaka, Japan, Cat# 168-23191) at 37 °C with 5% CO_2_ before the experiments.

### 6.6. Insulin secretion

Insulin secretion was measured using the static incubation method, as described previously (Murao et al., 2022; Murao et al., 2024).

To measure insulin secretion from cell lines, cells were seeded in 24-well plates at a density of 5 × 10^5^ cells/well and cultured for 48 h. On the day of measurement, the cells were subjected to three successive washes with 2.8G-KRBH, followed by a pre-incubation period of 30 min with 300 μL/well of 2.8G-KRBH. Subsequently, the supernatant was replaced with 300 μL/well of fresh KRBH containing the specified stimulations and incubated for 30 min at 37 °C. The reaction was terminated by cooling the plate on ice for ten minutes, after which the entire supernatant was collected for the quantification of released insulin using the homogeneous time-resolved fluorescence assay (HTRF) Insulin Ultrasensitive kit (Revvity, Waltham, MA, USA, Cat# 62IN2PEH) in accordance with the manufacturer’s instructions. Fluorescence was measured using an Infinite F Nano+ microplate reader (Tecan, Zürich, Switzerland).

To measure insulin secretion from islets, overnight cultured islets were rinsed twice with 2.8G-KRBH, followed by pre-incubation with 2.8G-KRBH for 30 min at 37 °C. Size-matched islets were hand-picked and dispensed in a 96-well plate (Corning, Glendale, AZ, USA, Cat# 353072) at 5 islets/well. KRBH (100 μL/well) containing the specified stimulations was added and incubated for 30 min at 37 °C. The supernatant was subjected to insulin quantification as described above.

### 6.7. Measurement of inositol 1-phosphate (IP1)

To measure IP1 content in cell lines, the cells were seeded in a 96-well plate (Corning, Glendale, AZ, USA, Cat# 353072) at a density of 1.0 × 10^5^ cells/well and cultured for 48h. On the day of measurement, the cells were subjected to three successive washes with IP1 assay buffer (146 mM NaCl, 4.2 mM KCl, 50 mM LiCl, 1 mM CaCl_2_, 0.5 mM MgCl_2,_ 10 mM HEPES, 0.1% BSA, pH7.4) supplemented with 2.8 mM glucose (2.8G-IP1 assay buffer), followed by a pre-incubation period of 30 min with 60 μL/well of 2.8G-IP1 assay buffer. Subsequently, 20 μL/well of IP1 assay buffer containing the specified stimulations at 4× concentration was added to the cells and incubated for another 30 min at 37 °C. The cells were lysed by the addition of 20 μL/well lysis buffer from the HTRF IP-One Gq Detection Kit (Revvity, Waltham, MA, USA, Cat# 62IPAPEB). 30 μL/well of the lysate were transferred to a 384-well low-volume plate (Cat #3826; Corning, Glendale, AZ, USA) and used for IP1 quantification according to the instructions of the HTRF IP-One Gq Detection Kit.

To measure IP1 content in mouse islets, overnight-cultured islets were rinsed twice with 2.8G-IP1 assay buffer, followed by pre-incubation with 2.8G-IP1 assay buffer for 30 min at 37 ℃. Size-matched islets were hand-picked and dispensed in a 96-well plate 96-well plate (Corning, Glendale, AZ, USA, Cat# 353072) at 5 islets/well. The islets were pre-incubated for 30 min with 45 μL/well of 2.8G-IP1 assay buffer. Subsequently, 15 μL/well of IP1 assay buffer containing the specified stimulations at 4× concentration was added to the cells and incubated for another 30 min at 37 °C. The cells were lysed by the addition of 15 μL/well of lysis buffer supplied in the kit. The lysate (30 μL/well) was subjected to IP1 quantification, as described above.

### 6.8. Imaging of intracellular Ca^2+^

Ca^2+^ imaging was performed as previously described (Murao et al., 2024). Briefly, cells were seeded in a 35 mm glass-bottom dish (Matsunami Glass, Osaka, Japan, Cat# D11530H) at a density of 1.28 × 10^5^ cells/dish and cultured for 48 h. Subsequently, the cells were loaded with 1 μM Fluo-4 AM (Dojindo, Kumamoto, Japan, Cat# F312) in 2.8G-KRBH for 20 min at 37 °C in room air. Following a brief washing, cells were loaded with 1 mL of fresh 2.8G-KRBH and basal recordings were performed for 300 s (from time -300 to 0). Immediately after the addition of 1 mL KRBH supplemented with stimulations at 2× concentration, recordings were resumed for another 600 s (from time 0 to 600) with a time interval of 2 s.

Time-lapse images were obtained using a Zeiss LSM 980 Airyscan2 inverted confocal laser scanning super-resolution microscope equipped with a Plan Apo 40×, 1.4 Oil DICII objective lens (Carl Zeiss Microscopy, Jena, Germany). The cells were excited at 488 nm laser with 0.3% output power, and fluorescence emission was measured at 508-579 nm. During observation, the cells were maintained at 37 °C using an incubator XLmulti S2 DARK (Pecon, Erbach, Germany).

Images were acquired in the frame mode at a rate of 2 frames per second and with an image size of 212.2 × 212.2 μm (512 × 512 pixels). The obtained images were analyzed using the ZEN 3.0 imaging software (Carl Zeiss Microscopy, Jena, Germany, RRID:SCR_021725). Cells were randomly chosen for analysis for each stimulation, and the number of cells analyzed is indicated in the figure legends. The fluorescence intensity of the entire cell body (F) was monitored and normalized to the average fluorescence intensity between -300 and 0 s (F0).

### 6.9. Imaging of intracellular ATP/ADP ratio^+^

The Cells were seeded in a 35 mm glass-bottom dish (Matsunami Glass, Osaka, Japan, Cat# D11530H) at a density of 6.0 × 10^4^ cells/dish and cultured for 48 h. Subsequently, the cells were transfected with 0.1 μg/dish of plasmid construct GW1-PercevalHR using Effectene transfection reagent (Qiagen, Venlo, Netherlands, Cat# 301425) in accordance with the manufacturer’s instructions and were cultured for another 48 h. On the day of the measurement, after a brief washing, the cells were loaded with 1 mL of fresh 2.8G-KRBH, and basal recordings were performed for 300 s (from time -300 to 0). Immediately after the addition of 1 mL KRBH supplemented with stimulations at 2× concentration, recordings were resumed for another 600 s (from time 0 to 600) with a time interval of 2 s.

The cells were excited at 445 nm and 488 nm laser with 0.8% and 0.3% output power, respectively, and fluorescence emission was measured at 508-543 nm. The ratio (R) of the fluorescence at 445 nm and 488 nm was calculated and normalized to the average fluorescence intensity between -300 and 0 s (R0). The other microscope configurations were the same as those used for Ca^2+^ imaging (Section 6.8).

### 6.10. Knockdown of Ketohexokinase (*Khk*) using small interfering RNA (siRNA)

siRNAs targeting *Khk* (Dharmacon, Lafayette, CO, USA, Cat# M-062217-01-0005) and non-targeting siRNA (Dharmacon, Lafayette, CO, USA, Cat# D-001206-14-50) were reverse-transfected using the DharmaFECT 2 transfection reagent (Dharmacon, Lafayette, CO, USA, Cat# T-2002-03). Briefly, a complex of siRNA and DharmaFECT 2 was prepared in serum-free DMEM (Sigma-Aldrich, St. Louis, MO, USA, Cat# D5796) at a volume of 100 μL/well according to the manufacturer’s instructions. Cells were resuspended in complete culture media at 1.25 × 10^6^ cells/mL. The cell suspension was then combined with siRNA/DharmaFECT 2 complex. The final concentrations of siRNA and DharmaFECT 2 were 40 nM and 0.4%, respectively. For insulin secretion and RT-qPCR experiments, the cells were seeded in 24-well plates (Corning, Glendale, AZ, USA, Cat# 353047) at 5 × 10^5^ cells/500 μL/well. For immunoblotting, the cells were seeded in 12-well plates (Corning, Glendale, AZ, USA, Cat# 353053) at 1 × 10^6^ cells/mL/well. For metabolic flux analysis, cells were seeded in 6-well plates (Cat #353046; Corning, Glendale, AZ, USA) at a density of 2 × 10^6^ cells/2 mL/well. The subsequent experiments were performed after a 48-hour culture.

### 6.11. Reverse transcription quantitative polymerase chain reaction (RT-qPCR)

cDNA was prepared using CellAmp Direct Lysis and RT set (Takara Bio, Shiga, Japan, Cat# 3737S/A) according to the manufacturer’s instructions. Quantitative real-time PCR was performed on a QuantStudio 7 Flex system (Thermo Fisher Scientific, Waltham, MA, USA, RRID:SCR_020245) using TaqMan Universal Master Mix II with UNG (Thermo Fisher Scientific, Waltham, MA, USA, Cat# 4440038) and Taqman probes: *Khk* (Cat# Mm00434647_m1) and TATA-box binding protein (*Tbp*, Cat# Mm01277042_m1). Relative gene expression of *Khk* was calculated using the 2^-ΔΔCT^ method and normalized to *Tbp*.

### 6.12. Immunoblotting

The cells were lysed with 50 μL/well of RIPA buffer (50 mM Tris-HCl pH 7.4, 150 mM NaCl, 1% NP-40, 0.25% sodium deoxycholate, 1% SDS), and 1x complete protease inhibitor cocktail (Sigma-Aldrich, St. Louis, MO, USA, Cat# 11697498001). The lysate was sonicated for 20 s on ice and centrifuged at 15000 × g at 4 °C for 10 min. The supernatant was collected and separated on a 7.5% polyacrylamide-SDS gel and transferred to a PVDF membrane. The membranes were blocked with 3% bovine serum albumin (BSA) (Sigma-Aldrich, St. Louis, MO, USA, Cat# A7906) in Tris-Buffered Saline with Tween 20 (TBS-T) and incubated with anti-KHK mouse monoclonal antibody (1:100) (Santa Cruz Biotechnology, Dallas, TX, USA, Cat# sc-377411) in TBS-T supplemented with 3% BSA overnight at 4 °C. The membrane was then incubated with HRP-conjugated anti-mouse immunoglobulins (1:2000) (Agilent, Santa Clara, CA, USA, Cat# P0447, RRID:AB_2617137) in TBS-T supplemented with 1% BSA for 1 h at room temperature. Signals were visualized using ECL Prime detection reagent (Cytiva, Buckinghamshire, UK, Cat# RPN2232). Images were taken using ImageQuant 800 (Cytiva, Buckinghamshire, UK). Subsequently, the membrane was stripped using Restore Western Blot Stripping Buffer (Thermo Fisher Scientific, Waltham, MA, USA, Cat# 21059) for 15 min and re-probed with an anti-α-tubulin mouse monoclonal antibody (1:1000) (Thermo Fisher Scientific, Waltham, MA, USA, Cat# A11126, RRID:AB_2534135). The images were quantified using ImageJ (version 1.53k, https://imagej.nih.gov/ij/index.html, RRID:SCR_003070).

### 6.13. Metabolic flux analysis

The cells were seeded in 6-well plates (Cat #353046; Corning, Glendale, AZ, USA) at 2 × 10^6^ cells/2 mL/well and cultured for 48h. The cells were then rinsed three times with 2.8G-KRBH, followed by pre-incubation with 2.8G-KRBH for 60 min at 37 °C. The supernatant was replaced with KRBH supplemented with specified stimulations including [U-^13^C]-glucose (Sigma-Aldrich, St. Louis, MO, USA, CAS: 110187-42-3 Cat# 660663) and [U-^13^C]-fructose (Cambridge Isotope Laboratories, Tewksbury, MA, USA, CAS: 287100-63-4, Cat# CLM-1553-1). The cells were incubated for 30 min at 37 °C and the supernatant was discarded. Cells were quickly rinsed with ice-cold water, extracted by the addition of 500 μL ice-cold extraction buffer (67.5% methanol, 7.5% chloroform, and 25% water), and the whole plate was snap-frozen in liquid nitrogen. Cells were thawed on ice, scraped into 2 mL screw tubes along with the supernatant, and stored at -80 °C until extraction of the metabolites.

For extraction of the metabolites, samples were supplemented with 80 μL of diluted (1:640) internal standard (Human Metabolome Technologies, Yamagata, Japan, Cat# H3304-1002), 165 μL of methanol, and 465 μL of chloroform. The samples were then homogenized using a pre-cooled bead crusher at 3200 rpm for 1 min and centrifuged at 15000 × g at 4 °C for 3 min. The aqueous layer was transferred to pre-wetted ultrafiltration tubes (Human Metabolome Technologies, Yamagata, Japan, Cat# UFC3LCCNB-HMT) and centrifuged at 9100 × g, 4 °C until completely filtrated. The filtrate was freeze-dried, re-dissolved in 10 μL of water, and subjected to mass spectrometry. The organic layer was evaporated by decompression at room temperature and the residue was resuspended in lysis buffer (see Section 6.7), which was then subjected to a BCA protein assay (Thermo Fisher Scientific, Waltham, MA, USA, Cat# 23225).

cGMP was measured by G7100A capillary electrophoresis (Agilent Technologies, Santa Clara, CA, USA) interfaced with a G6224A time-of-flight LC/MS mass spectrometer (Agilent Technologies, Santa Clara, CA, USA). A G1310A isocratic pump (Agilent Technologies, Santa Clara, CA, USA) equipped with a G1379B degasser (Agilent Technologies, Santa Clara, CA, USA) was used to supply sheath liquid (Human Metabolome Technologies, Yamagata, Japan, Cat# H3301-2020). The mass spectrometer was operated in the negative ionization mode. All separations were performed on fused silica capillaries (Human Metabolome Technologies, Yamagata, Japan, Cat# H3305-2002) at 25 °C using an anion analysis buffer (Human Metabolome Technologies, Yamagata, Japan, Cat# H3302-2021) as the background electrolyte. The applied voltage was set to 30 kV at 20 °C, together with a pressure of 15 mbar. Sheath liquid was delivered to a nebulizer by an isocratic pump at 1 mL/min. Chromatograms and mass spectra were analyzed by MassHunter qualitative analysis version 10.0 (Agilent Technologies, Santa Clara, CA, USA, RRID:SCR_015040). Annotation and quantification of chromatogram peaks were carried out using a standard mixture (Human Metabolome Technologies, Yamagata, Japan, Cat# H3302-2021) as a reference.

### 6.14. Statistical Analysis

Sample sizes were estimated from the expected effect size based on previous experiments. No randomization or blinding was used. For cell line experiments, including insulin secretion, IP1 measurement, RT-qPCR, immunoblotting, and metabolic tracing, *n* represents the number of biological replicates of cells grown in different wells of the same multiwell plate. For islet experiments, including insulin secretion and IP1 measurement, *n* represents the number of wells, each containing five islets. For Ca^2+^ and ATP/ADP ratio measurements, *n* represents the number of different single cells analyzed. Data are shown as the mean ± standard deviation (SD) along with the plot of individual data points. For statistical comparisons between two groups, a two-tailed unpaired Welch’s unpaired *t*-test was used. For comparisons between three groups, Welch’s one-way analysis of variance (ANOVA) was followed by pairwise comparisons corrected using Dunnett’s method. Normality of the distribution was confirmed by the Shapiro-Wilk test. P-values less than 0.05 were considered statistically significant and are indicated in the figures. P-values more than 0.05 are not indicated in the figures. The statistical analyses used are indicated in the figure legends. Statistical analyses were performed using GraphPad Prism 10 (Graphpad Software, Boston, MA, USA, https://www.graphpad.com; RRID:SCR_002798).

## 7 RESULTS

### 7.1 FSIS is dependent on K_ATP_ channels and calcium influx

We first examined the responsiveness of B6 mouse islets and MIN6-K8 cells to various doses of fructose. Fructose, at concentrations ranging from 2.5 mM to 12.5 mM and 0.3 mM to 10 mM, dose-dependently increased insulin secretion in B6 mouse islets (Figure 1A) and MIN6-K8 cells (Figure 1B), respectively, when glucose levels were at a stimulatory level (8.8 mM). However, fructose displayed no effect on insulin secretion at basal glucose level (2.8 mM) in MIN6-K8 cells (Figure 1C), indicating that FSIS is glucose-dependent.

**Figure 1.**
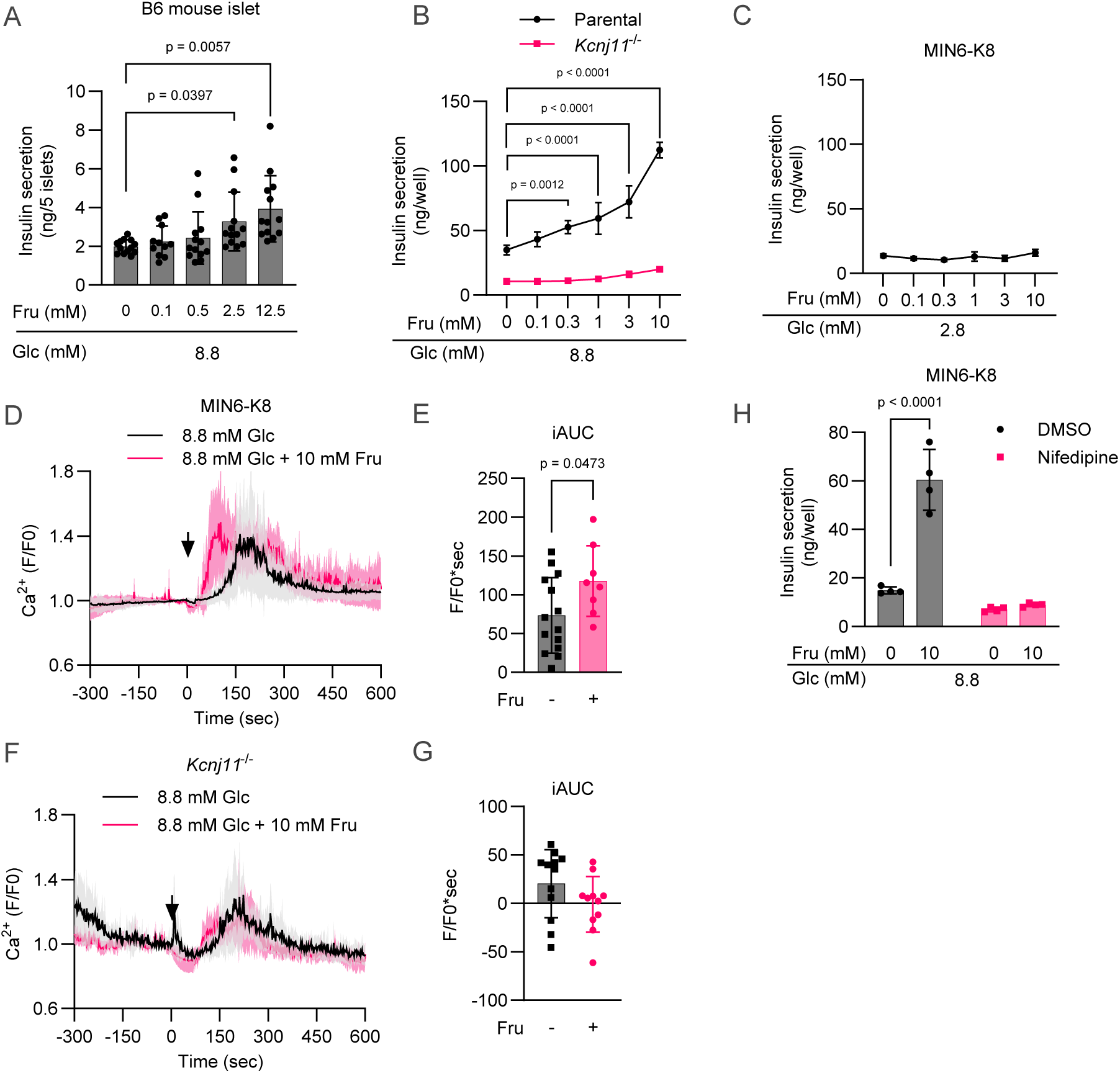
FSIS is dependent on K_ATP_ channels and calcium influx. (A) Dose-dependent effect of fructose on insulin secretion at 8.8 mM glucose in B6 mouse islets. n = 11–13. (B) Dose-dependent effect of fructose on insulin secretion at 8.8 mM glucose in parental MIN6-K8 and *Kcnj11*^-/-^ cells. n = 4. (C) Effect of fructose on insulin secretion at 2.8 mM glucose in MIN6-K8 cells. n = 4. (D) –(G) Effect of fructose on intracellular Ca^2+^ levels measured using Fluo-4. The time course of normalized fluorescence intensity at 508-579 nm is indicated in (D) and (F). Black arrows represent the addition of the indicated stimulations at time = 0. The magnitude of Ca^2+^ responses was quantified as iAUC in (E) and (G), using F/F0 =1 as the baseline. (D)–(E): MIN6-K8 cells. 8.8 mM Glc: n = 14; 8.8 mM Glc + 10 mM Fru: n = 8. (F)–(G): *Kcnj11*^-/-^ cells. 8.8 mM Glc: n = 12; 8.8 mM Glc + 10 mM Fru: n = 11. (H) Effect of nifedipine (10 μM) on FSIS in MIN6-K8 cells. n = 4. Data are presented as the mean ± standard deviation (SD). Glc, glucose; Fru, fructose. Statistical comparisons were performed using Welch’s two-way ANOVA with Dunnett’s post-hoc test for (A) and (B), and Welch’s unpaired two-tailed t-test for (E), (G), and (H).

To explore whether intracellular fructose metabolism is coupled with insulin secretion in a manner comparable to that of glucose, the involvement of K_ATP_ channels and intracellular Ca^2+^ in FSIS was examined. We utilized *Kcnj11*^-/-^ β-cells, which lack K_ATP_ channel activity and exhibit continuous cell membrane depolarization regardless of glucose metabolism (Oduori et al., 2020). Fructose did not significantly increase insulin secretion in *Kcnj11*^-/-^ β-cells at 8.8 mM glucose (Figure 1B), suggesting that FSIS is dependent on K_ATP_ channel activity. This finding is supported by the previous observation that FSIS was abolished by clamping the K_ATP_ channel in the open state with diazoxide in MIN6-K8 cells (Seino et al., 2015).

In MIN6-K8 cells, fructose augmented the increase in intracellular Ca^2+^ ([Ca^2+^]i) levels induced by 8.8 mM glucose (Figure 1, D–E). In *Kcnj11*^-/-^ β-cells, fructose did not increase [Ca^2+^]i levels (Figure 1, F–G). These trends in [Ca^2+^]i responses were consistent with the differential FSIS observed in Figure 1B. Furthermore, Nifedipine, an inhibitor of L-type VDCCs, abolished FSIS in MIN6-K8 cells (Figure 1H). Overall, these findings indicate that K_ATP_ channels and Ca^2+^ influx are necessary for FSIS.

### 7.2 Characterization of β-cell fructose metabolism

According to our previously deposited RNA-seq datasets (Hashim et al., 2018), *Slc2a5* (GLUT5) and *Khk* are expressed in both MIN6-K8 cells and B6 mouse islets (Supplementary Table 1). To determine whether fructose is metabolized in β-cells, we traced the metabolic fate of ^13^C-labelled fructose. MIN6-K8 cells were stimulated for 30 min with 10 mM [U-^13^C]-fructose, a stable isotopomer of fructose in which all six carbon atoms have been replaced with ^13^C, in the presence of 8.8 mM unlabeled glucose. For each intermediate, ^13^C-labelled isotopomers were quantified by capillary electrophoresis-time-of-flight mass spectrometry (CE-TOFMS) (Figure 2). M–M6 represents the isotopomers in which 0–6 carbon atoms were replaced with ^13^C. Naturally occurring isotopomers are mostly M and M1, whereas incorporation of [U-^13^C]-fructose-derived carbons generates M2–M6.

**Figure 2.**
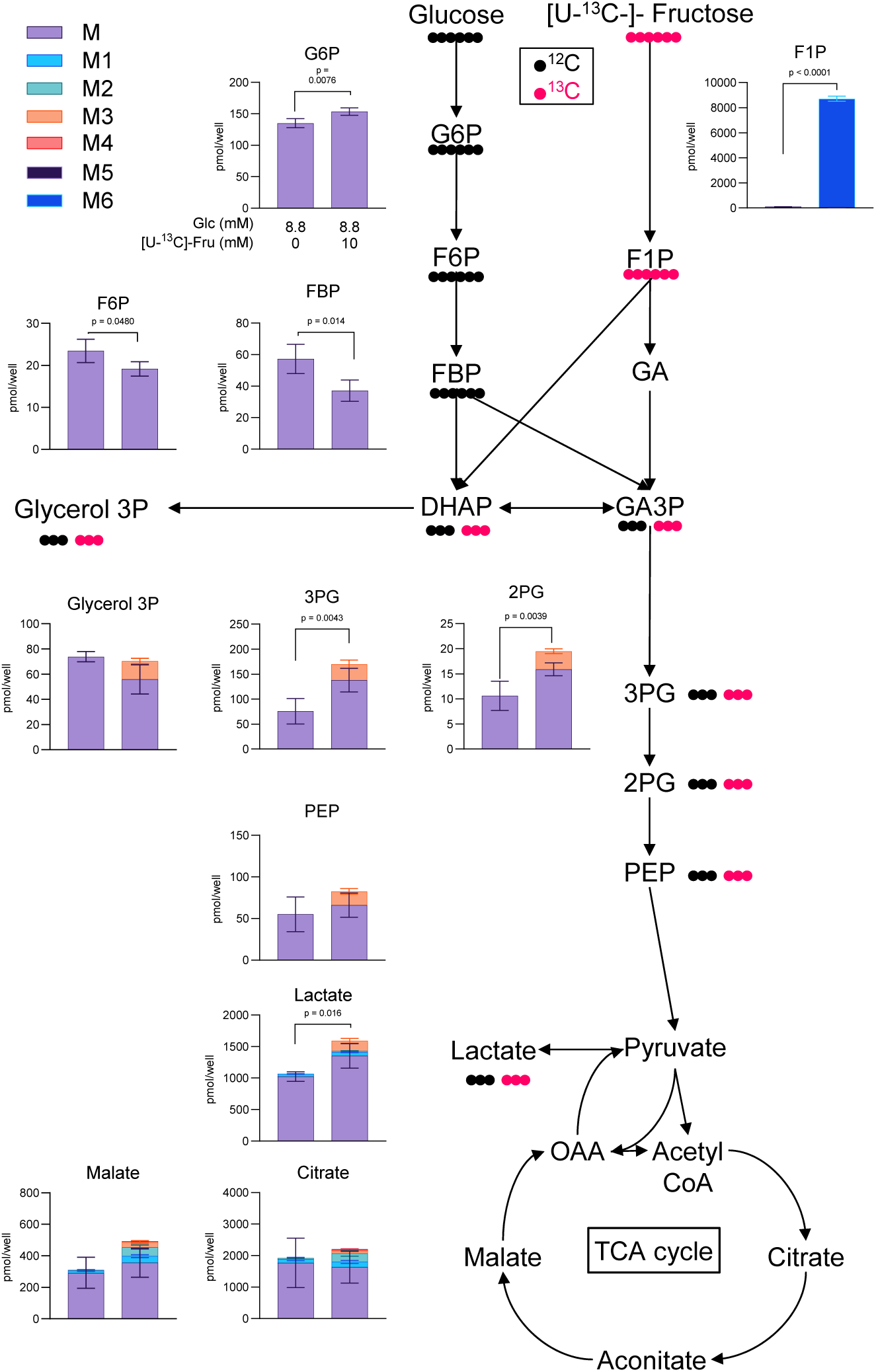
[U-^13^C]-fructose tracing in MIN6-K8 cells. The total content and isotopic distribution of each intermediate are indicated by stacked bar graphs alongside the schematic overview of metabolic fates of glucose and [U-^13^C]-fructose. n = 4. Data are presented as the mean ± standard deviation (SD). Statistical comparisons were made between the total content using Welch’s unpaired two-tailed t-test.

Upper glycolytic intermediates, such as G6P, F6P, and FBP, did not comprise M2–M6, whereas F1P was found only as M6 under [U-^13^C]-fructose treatment. Lower glycolytic intermediates such as 3PG, 2PG, PEP, glycerol 3P, and lactate were labelled as M3. Meanwhile, TCA cycle intermediates such as citrate and malate were labelled as M2-M6. These results indicate that [U-^13^C]-fructose enters lower glycolysis and the subsequent TCA cycle. We also assessed labeling efficiency by ^13^C fraction of the total carbon atoms in each intermediate (Supplementary Figure 1A). ^13^C Fraction was negligible at the basal state and increased up to 0.1–0.2 in the presence of [U-^13^C]-fructose in glycolysis and TCA cycle intermediates.

Because FSIS was glucose-dependent (Figure 1, B–C), we sought to clarify whether fructose affects glucose metabolism. To this end, MIN6-K8 cells were labelled with 10 mM unlabeled fructose in the presence of 8.8 mM [U-^13^C]-glucose (Figure 3A). Glucose-derived ^13^C was robustly incorporated into all intermediates, except for F1P. The upper and lower glycolytic intermediates were labelled as M6 and M3, respectively. In those intermediates, the ^13^C fraction was 0.3–0.5, a value greater than that observed using [U-^13^C]-fructose at a higher concentration (10 mM), suggesting that the rate of fructose metabolism is lower than that of glucose in β-cells (compare Figure 3B and Supplementary Figure 1A).

**Figure 3.**
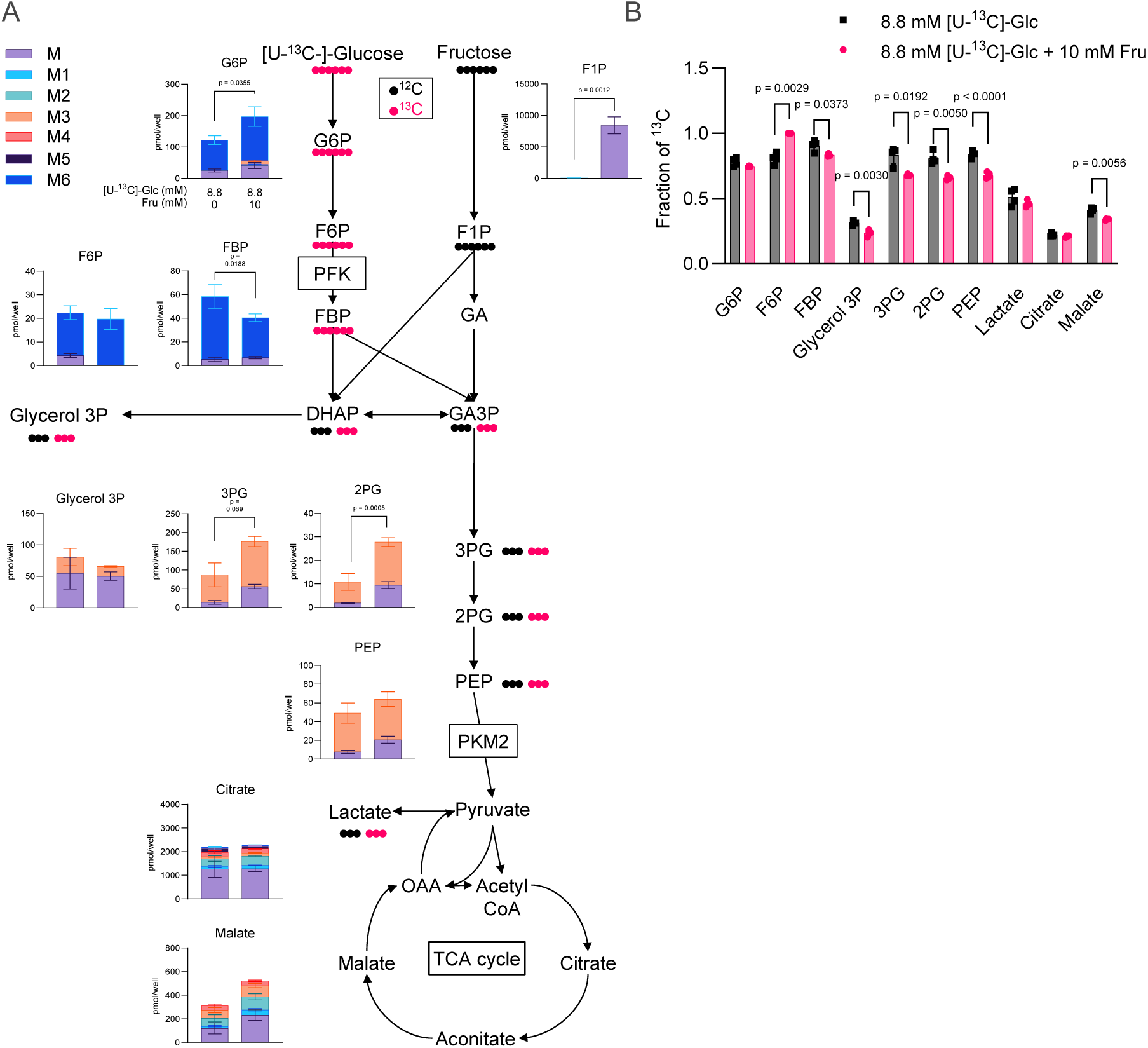
[U-^13^C]-glucose tracing in MIN6-K8 cells. (A) The total content and isotopic distribution of each intermediate are indicated as stacked bar graphs alongside a schematic overview of the metabolic fates of [U-^13^C]-glucose and fructose. n = 4. Statistical comparisons were made between the total content using Welch’s unpaired two-tailed t-test. (B) Fraction of ^13^C in each intermediate. Statistical comparisons were made using Welch’s unpaired two-tailed t-test. Data are presented as the mean ± standard deviation (SD).

Notably, the content (total M–M6) and labeling of several intermediates were significantly affected by fructose. The contents of G6P, 3PG, and 2PG were increased, while that of FBP was decreased by fructose, a trend consistent regardless of the ^13^C source (Figure 3A and Figure 2). Additionally, glucose-derived ^13^C fraction was increased in FBP but was decreased in later steps of glycolysis (Figure 3B). Previous studies have shown that high concentrations of fructose inhibit phosphofructokinase (PFK) (Kreuzberg, 1978) and pyruvate kinase (PK) (Mapungwana and Davies, 1982; Taylor et al., 2021). As both PFK and PK reactions are irreversible (Chandel, 2021), it is reasonable to assume that the reduction in their activities causes the accumulation of upstream intermediates and/or the depletion of downstream intermediates. Therefore, our interpretation was that (1) increased content and labeling of G6P, decreased content of FBP, and decreased labeling of intermediates after FBP (Figure 3, A–B) reflect reduced PFK activity, and that (2) increased contents of 3PG and 2PG are due to reduced PK activity. Overall, our metabolic tracing revealed interactions between fructose metabolism and glycolysis, indicating that fructose enters lower glycolysis but simultaneously restricts glycolysis by inhibiting PFK and PK.

### 7.3 β-cell fructose metabolism has little effect on insulin secretion

We next investigated the impact of intracellular fructose metabolism on cellular energy status in MIN6-K8 cells. Fructose did not significantly alter the levels of adenine nucleotides (Supplementary Figure 1, B–D) or redox cofactors (NAD, NADH, NADP, and NADPH) (Supplementary Figure 1, F–I). However, the ATP/ADP ratio tended to decrease (Supplementary Figure 1E). Indeed, time-lapse imaging using PercevalHR, a genetically encoded fluorescent biosensor (Tantama et al., 2013), revealed that fructose suppressed the glucose-induced increase in the ATP/ADP ratio (Figure 4, A–B).

**Figure 4.**
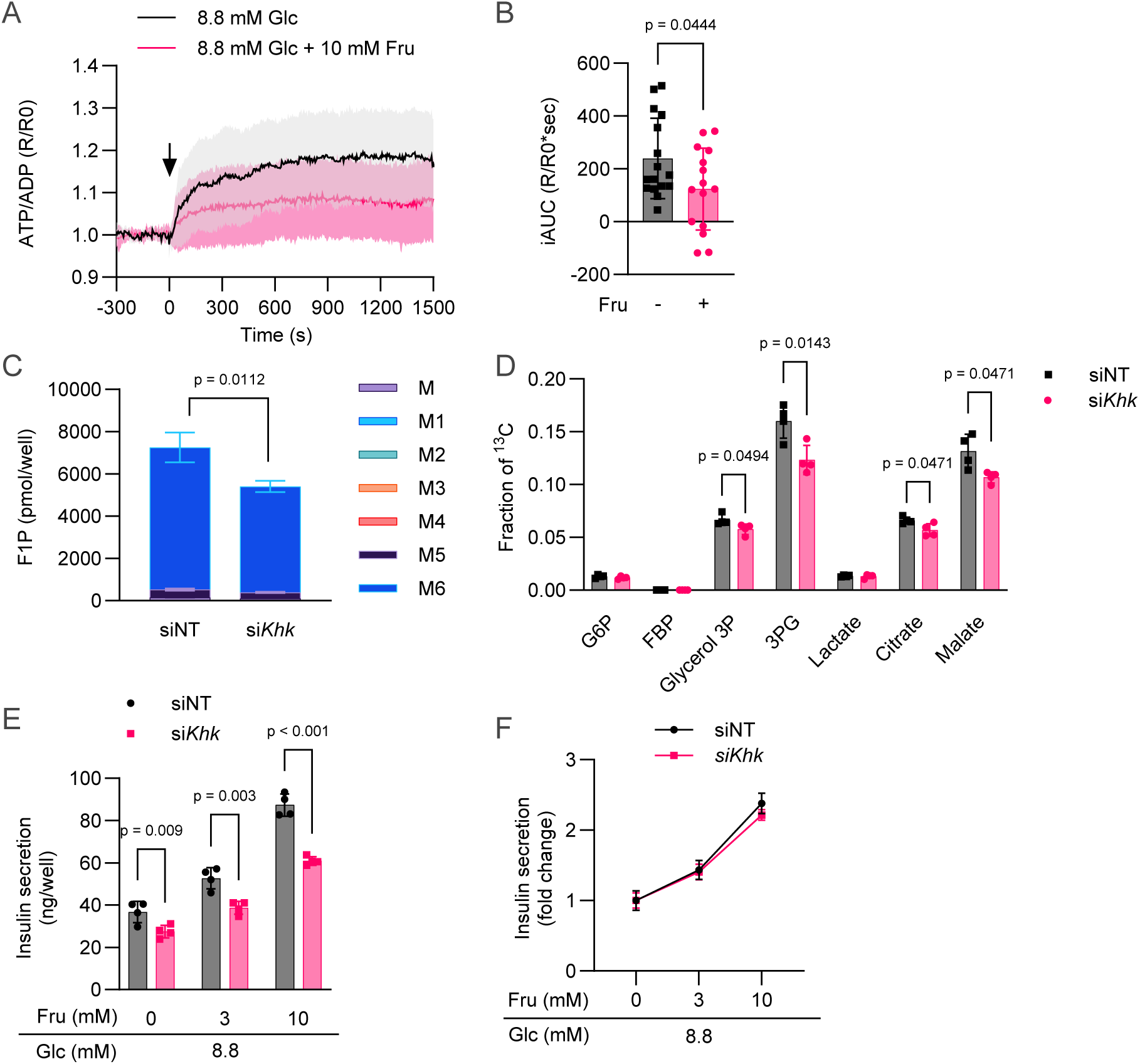
Suppression of intracellular fructose metabolism does not affect fructose responsiveness in MIN6-K8 cells. (A) –(B) Effect of fructose on intracellular ATP/ADP ratio in MIN6-K8 cells as visualized by Perceval-HR. 8.8 mM Glc: n = 16, 8.8 mM Glc + 10 mM Fru: n=15. (A) Time course of normalized ratiometric signals (R/R0). (B) The magnitude of the responses was quantified as iAUC using R/R0 = 1 as the baseline. (E) –(D) [U-^13^C]-fructose tracing under Khk knockdown in MIN6-K8 cells. The total content and isotopic distribution of F1P are presented in (C). The fraction of ^13^C in each intermediate is presented in (D). n = 4. See Supplementary Figure 2 for the total content of other metabolites. (F) The effect of Khk knockdown on FSIS was expressed as fold change over 0 mM fructose. See Supplementary Figure 3D for the original values. All experiments were performed using MIN6-K8 cells. siNT, non-targeting siRNA. Data are presented as the mean ± SD. Statistical comparisons were performed using Welch’s unpaired two-tailed t-test.

To clarify how intracellular fructose metabolism affects FSIS, we depleted KHK using siRNA. KHK mRNA and protein levels were reduced by approximately 70% and 20%, respectively (Supplementary Figure 2, A–C). Since the knockdown efficiency was moderate at the protein level, we also validated the metabolic phenotypes using [U-^13^C]-fructose tracing. As expected, KHK knockdown decreased M6 F1P (Figure 4C). Although total content of other intermediates remained unchanged (Supplementary Figure 3), ^13^C fraction in later glycolysis and TCA intermediates decreased (Figure 4D), confirming that KHK knockdown significantly suppressed intracellular fructose metabolism. Knockdown of KHK resulted in a small decrease in insulin secretion, regardless of the presence of fructose (Figure 4E). However, when expressed as the fold-change over 0 mM fructose, KHK knockdown had no effect on fructose responsiveness (Figure 4F). These results provide evidence against the hypothesis that β-cell fructose metabolism fuels insulin secretion via ATP production.

#### β-cell fructose responsiveness parallels PLC signaling

Given the limited role of fructose metabolism in FSIS, we investigated the molecular basis of β-cell fructose responsiveness.

Fructose functions as a substrate for the sweet taste receptor (STR), a G protein-coupled receptor complex composed of taste receptor type 1 member 2 (TAS1R2) and 3 (TAS1R3) (Nelson et al., 2001; Zhao et al., 2003; Chandrashekar et al., 2006). STR signals through a second messenger system that involves G-protein gustducin, phospholipase C β2 (PLCβ2), inositol 1,4,5-trisphosphate (IP_3_), and transient receptor potential M4/5 (TRPM4/5) (Huang et al., 1999; Zhang et al., 2007). This pathway was originally implicated in taste transduction in the tongue (Ahmad and Dalziel, 2020). Later studies have shown that STRs are expressed in β-cells (Nakagawa et al., 2009) and regulate FSIS (Kyriazis et al., 2012). Indeed, U73122, an inhibitor of PLCβ2 (Bleasdale et al., 1990), partially inhibited FSIS in MIN6-K8 cells (Figure 5A).

**Figure 5.**
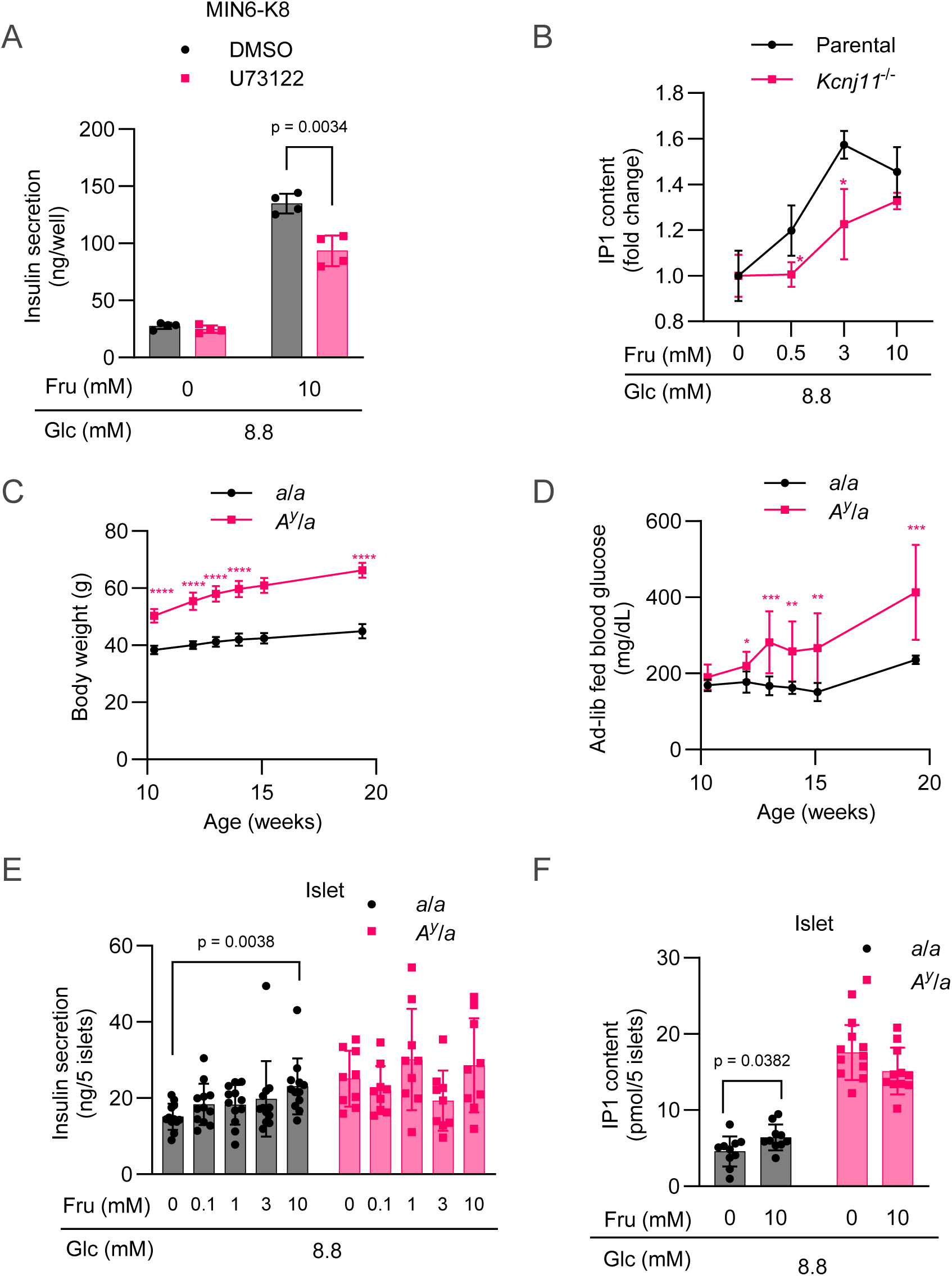
Association between β-cell fructose responsiveness and PLC signaling. (A) Effect of U73122 (20 μM) on FSIS in MIN6-K8 cells. n = 4. (B) Dose-dependent effects of fructose on intracellular IP1 levels in parental MIN6-K8 and *Kcnj11*^-/-^ cells. The data are expressed as fold-change over 0 mM fructose. n = 6. The two cell lines were compared for statistical significance. *p < 0.05. (C) Age-dependent changes in body weight in NSY.B6 strains. The two strains were compared for statistical significance. *a*/*a*: n = 6; *A^y^*/*a*: n = 12. ****p < 0.0001. (D) Age-dependent changes in ad-lib fed blood glucose levels in NSY.B6 strains. The two strains were compared for statistical significance. *a*/*a*: n = 6; *A^y^*/*a*: n = 12. *p < 0.05, **p < 0.01, ***p < 0.001. (E) FSIS in islets isolated from NSY.B6 strains. *a*/*a*: n = 9; *A^y^*/*a*: n = 12. (F) Effects of fructose on intracellular IP1 levels in islets isolated from NSY.B6 strains. *a*/*a*: n = 10; *A^y^*/*a*: n = 11. Data are presented as the mean ± SD. Statistical comparisons were performed using Welch’s unpaired two-tailed t-test.

We sought to determine whether differential PLC activity can account for fructose responsiveness in various β-cell models. To assess fructose-induced PLC activity, we measured d-myo-inositol 1-phosphate (IP1) as a surrogate for IP_3_. 0.5 mM and 3 mM fructose caused a dose-dependent accumulation of IP1 in parental MIN6-K8 cells, but this increase was diminished in *Kcnj11*^-/-^ cells, suggesting that diminished PLC signaling underlies impaired fructose responsiveness (Figure 5B).

The pathophysiological relevance of these findings was examined using diabetes mouse models. NSY.B6-*Tyr*^+^, *A^y^* is a spontaneous type 2 diabetes mouse strain that was recently established by crossing the NSY and B6J-*A^y^* strains, thereby interrogating the obesity-related agouti-yellow (*A^y^*) mutation in the agouti (*a*) gene into a diabetes-prone NSY background (Ohno et al., 2022). Consistent with the observation by Ohno et al., Ay heterozygous male mice (*A^y^*/*a*) displayed obesity and hyperglycemia after 13 weeks of age compared to wild-type male mice (*a*/*a*) (Figure 5, C–D). Islets were isolated at 19–20 weeks of age and tested for fructose responsiveness. Insulin secretion and IP1 levels in the a/a islets exhibited a significant increase at 10 mM fructose (Figure 5, E–F). In contrast, insulin secretion and IP1 levels in *A^y^*/*a* islets failed to respond to fructose. Hence, decreased fructose responsiveness in *Kcnj11*^-/-^ cells and diabetic islets correlated with impaired PLC activity. These findings suggest that β-cell fructose responsiveness is not dictated by intracellular fructose metabolism, but rather by STR-mediated PLC signaling.

### 7.4 PK activator counteracts fructose-mediated PK inhibition in β-cells

Improving the responsiveness of β-cells to fructose is of therapeutic importance. We hypothesized that neutralizing the inhibitory impact of fructose on glycolysis in β-cells would allow for the effective utilization of fructose as a fuel source for insulin secretion.

It has recently been shown that F1P binds to and inhibits the M2 isoform of pyruvate kinase (PKM2) in intestinal epithelial cells (Taylor et al., 2021). Notably, this inhibition can be overcome by TEPP-46, a small-molecule PK activator (Jiang et al., 2010). Considering that PKM2 (*Pkm*) is the primary PK isoform in MIN6-K8 cells and B6 mouse islets, as evidenced by our previously deposited RNA-seq datasets (Supplementary Table 1) (Hashim et al., 2018), this mechanism is also likely to be applicable to our study.

Motivated by these findings, we sought to determine whether TEPP-46 enhanced β-cell fructose responsiveness. The effect of TEPP-46 on PK activity was evaluated using lysates from MIN6-K8 cells. TEPP-46 significantly elevated PK activity, regardless of whether the cells were pretreated with fructose (Figure 6A). However, fructose pretreatment caused only a slight decrease in PK activity, implying that the binding of intracellular F1P to PK is disrupted by cell lysis.

**Figure 6.**
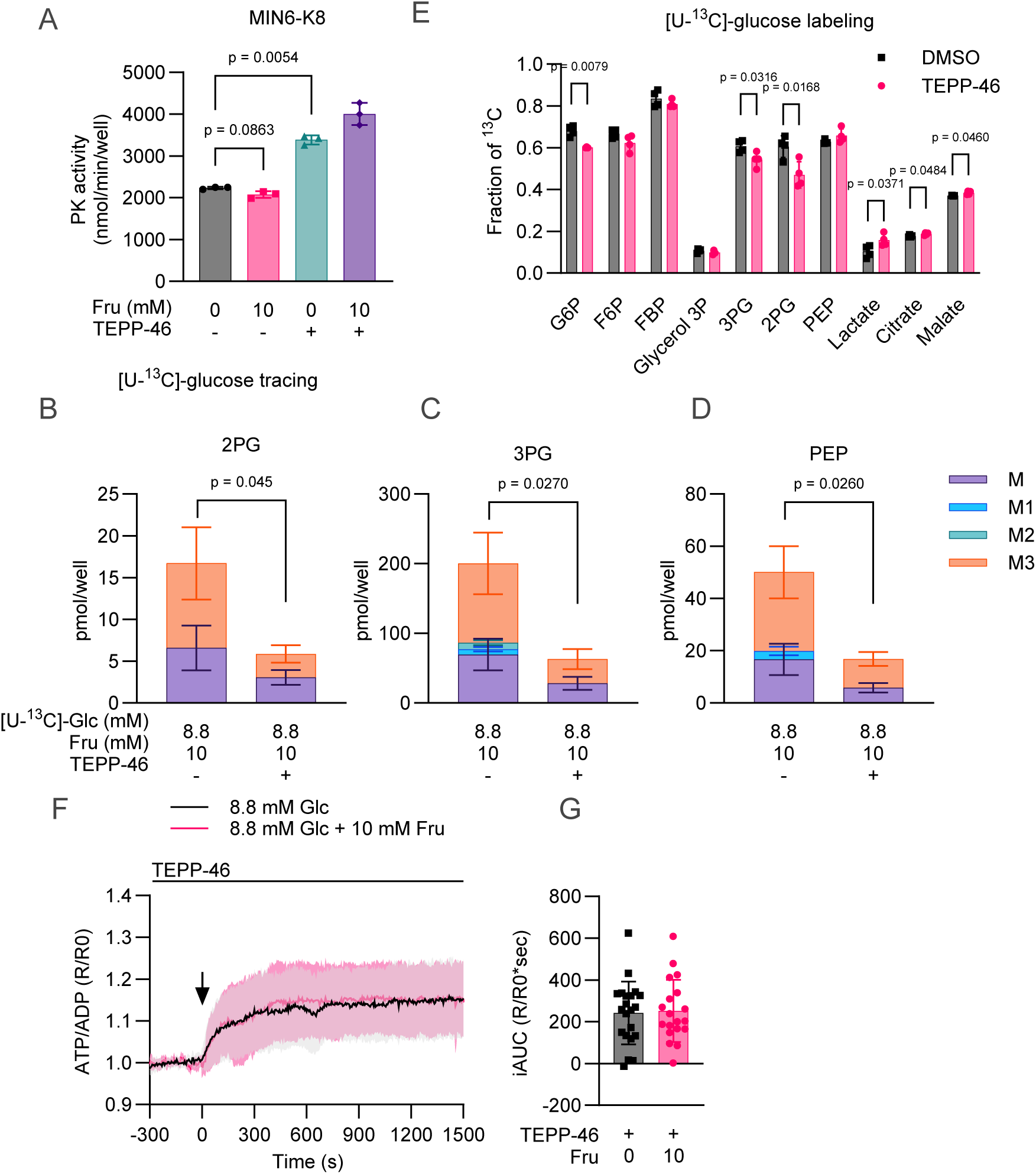
PK activator counteracts fructose-mediated PK inhibition in β-cells. (A) Effect of TEPP-46 on PK activity in MIN6-K8 cell lysates. n = 3. (B) –(E) [U-^13^C]-glucose tracing after TEPP-46 treatment. The total content and isotopic distribution of the lower glycolytic intermediates are presented in (B)–(D). The fraction of ^13^C in each intermediate is presented in (E). n = 4. See Supplementary Figure 4 for the total content and isotopic distribution of the other metabolites. (F) –(G) Effect of fructose on intracellular ATP/ADP ratio in the presence of TEPP-46 as visualized by Perceval-HR. 8.8 mM Glc: n = 21; 8.8 mM Glc + 10 mM Fru: n = 19. (F) Time course of normalized ratiometric signals (R/R0). (G) The magnitude of the responses was quantified as iAUC using R/R0 = 1 as the baseline. All experiments were performed using MIN6-K8 cells. TEPP-46 was used at a concentration of 10 μM. Data are presented as the mean ± SD. Statistical comparisons were performed using Welch’s one-way ANOVA with Dunnett’s post-hoc test for (A) and Welch’s unpaired two-tailed t-test for (B) –(E).

Therefore, to confirm TEPP-46’s effect in situ, we conducted metabolic tracing experiments using either [U-^13^C]-glucose or [U-^13^C]-fructose in the presence of TEPP-46. In both experiments, fructose-induced accumulation of 2PG, 3PG, and PEP was substantially attenuated by TEPP-46 (Figure 6, B–D, Supplementary Figures 4 and 5A), indicating effective PK activation, as previously reported in hepatocytes (Abulizi et al., 2020).

In [U-^13^C]-glucose tracing, ^13^C fraction in 3PG and 2PG decreased, while that in later intermediates (lactate, citrate, and malate) increased (Figure 6E). Similarly, in [U-^13^C]-fructose tracing, the ^13^C fraction in citrate and malate slightly, but significantly, increased (Supplementary Figure 5B). These results suggest that TEPP-46 facilitates the metabolic flux of glucose and fructose after PK. Moreover, in the presence of TEPP-46, fructose did not alter glucose-induced increases in the ATP/ADP ratio, as monitored by Perceval HR (Figure 6, F– G). These results indicate that TEPP-46 was able to overcome fructose-mediated PK inhibition and counteract fructose-induced ATP depletion.

### 7.5 PK activator enhances fructose responsiveness in β-cells

TEPP-46 had no effect on insulin secretion at 8.8 mM alone, but it significantly increased insulin secretion when additional 10 mM fructose was present (Figure 7A). Consistently, TEPP-46 did not significantly alter the Ca^2+^ response to glucose alone (Figure 7, B–C), but augmented the increase in Ca^2+^ at 10 mM fructose (Figure 7, D–E). These findings indicate that TEPP-46 requires fructose to exert insulinotropic effects. KHK knockdown markedly suppressed fructose responsiveness in the presence of TEPP-46 (Figure 7F). Taken together, these observations suggest that TEPP-46 enables intracellular fructose metabolism to effectively fuel insulin secretion.

**Figure 7.**
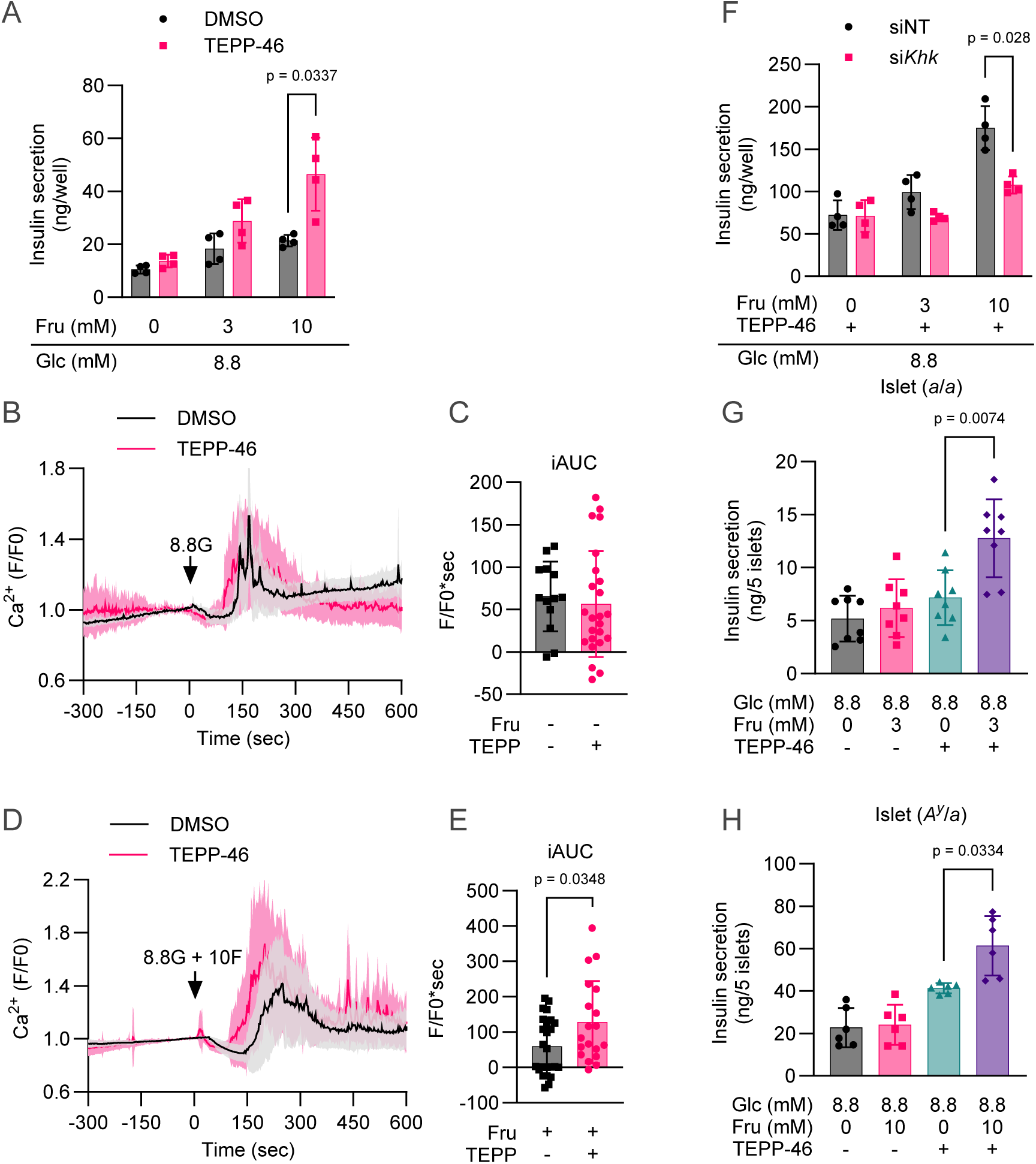
PK activator enhances β-cell fructose responsiveness. (A) Effect of TEPP-46 on FSIS in MIN6-K8 cells. n = 4. (B) (C) Effect of TEPP-46 on glucose-induced Ca^2+^ increase measured using Fluo-4. DMSO: n = 13; TEPP-46: n = 24. (B) The black arrow represents the addition of 8.8 mM glucose (8.8G) at time 0. (C) The magnitude of the responses was quantified as iAUC using F/F0 = 1 as the baseline. (C) –(E) Effect of TEPP-46 on fructose-induced Ca^2+^ increase measured using Fluo-4. DMSO: n = 25; TEPP-46: n = 24. (D) The black arrow represents the addition of 8.8 mM glucose supplemented with 10 mM fructose (8.8G + 10F) at time 0. (E) The magnitude of the responses was quantified as iAUC using F/F0 = 1 as the baseline. (F) Effect of Khk knockdown on FSIS in the presence of TEPP-46 in MIN6-K8 cells. siNT, non-targeting siRNA. n = 4. (G) Effect of TEPP-46 on FSIS in *a*/*a* mouse islets. n = 8. (H) Effect of TEPP-46 on FSIS in *A^y^*/*a* mouse islets. n = 6. Experiments (A)–(F) were performed using MIN6-K8 cells. TEPP-46 was used at a concentration of 10 μM. Data are presented as the mean ± SD. Statistical comparisons were performed using Welch’s unpaired two-tailed t-test for (A) –(E) and Welch’s one-way ANOVA with Dunnett’s post-hoc test for (G) –(H).

The effects of TEPP-46 were validated in both normal and diabetic islets. *a*/*a* mouse islets failed to display significant FSIS to 3 mM fructose alone, but showed marked FSIS when treated with TEPP-46 (Figure 7G). Similarly, *A^y^*/*a* islets were responsive to 10 mM fructose in the presence of TEPP-46 (Figure 7H). These results corroborate the effectiveness of PK activation in enhancing fructose responsiveness in normal and diabetic β-cells.

## 7 DISCUSSION

The present study demonstrated that (1) the responsiveness of β-cells to fructose is not regulated by intracellular fructose metabolism, but rather by PLC signaling; (2) fructose limits glycolysis by inhibiting PK and decreasing the ATP/ADP ratio; and (3) a PK activator overcomes the inhibition of PK caused by F1P, thereby enhancing β-cell responsiveness to fructose. These findings highlight the distinction between FSIS and GSIS, most of which is attributable to the unique metabolic property of fructose.

Fructose has been known to be metabolized in β-cells (Ashcroft et al., 1972; Zawalich et al., 1977; Sener and Malaisse, 1988; Giroix et al., 1999). However, the extent to which β-cell fructose metabolism contributes to insulin secretion has remained undetermined. In this study, by employing ^13^C-labelled metabolic tracing and Khk knockdown experiments, we sought to revisit this question.

In accordance with previous studies (Ashcroft et al., 1970; Zawalich et al., 1977; Sener and Malaisse, 1988; Giroix et al., 1999), we found that the rate of fructose metabolism was significantly lower than that of glucose metabolism in β-cells. Nevertheless, some studies have proposed that FSIS may be regulated by ATP produced through intracellular fructose metabolism, as the rates of insulin secretion and fructose metabolism or ATP production have been observed to be correlated (Zawalich et al., 1977; Sener and Malaisse, 1988). However, these findings have been challenged by later studies using diabetic GK rats (Giroix et al., 1999) and an INS-1E β-cell line (Bartley et al., 2019), which showed no impact of acute fructose on mitochondrial respiration or ATP production, as well as by our observation that fructose impedes the glucose-induced increase in the ATP/ADP ratio.

The relationship between intracellular fructose metabolism and ATP levels is complex and involves multiple factors. This study identified two factors that likely contribute to fructose-induced reduction in the ATP/ADP ratio: ATP consumption caused by KHK and fructose-mediated inhibition of PFK and PK. Phosphorylation of fructose by KHK requires ATP hydrolysis, which leads to a reduction in ATP levels in the liver and kidney cells (Van den Berghe et al., 1977; Abdelmalek et al., 2012; Cirillo et al., 2009). Chronic fructose treatment has also been shown to decrease ATP levels in INS-1E cells (Bartley et al., 2019).

In MIN6-K8 cells, M6 F1P accumulated substantially after [U-^13^C]-fructose treatment (approximately 40 times more than G6P), but ^13^C incorporation in downstream intermediates was relatively low. This might be due to the imbalance between the expression levels of GLUT5 and KHK and that of aldorase B, which was very low (Supplementary Table 1). Hence, we assume that ATP consumption by KHK cannot be balanced by subsequent ATP production resulting from fructose catabolism, leading to a decrease in the ATP/ADP ratio. Consequently, fructose catabolism and cellular energy status were not positively correlated.

Previous research has utilized radioisotope-labelled sugars to measure the utilization or oxidation rates of the sugar in the entire islet (Ashcroft et al., 1970; Zawalich et al., 1977; Sener and Malaisse, 1988; Giroix et al., 1999), reporting that glucose utilization rate was unaffected by the addition of excess fructose (Zawalich et al., 1977). In contrast, the present study observed an apparent decrease in glycolytic flux by fructose. These discrepant results may imply that the whole-cell glucose utilization rate is largely determined by glucokinase activity and does not necessarily reflect fluctuations in downstream steps. In this regard, the utilization of mass spectrometry with ^13^C-labelled sugars, which enabled direct tracing of carbon flux in each intermediate, offered a significant advantage in revealing the interactions between glycolysis and fructolysis.

In the liver, fructose promotes glucose uptake and phosphorylation by stimulating the release of glucokinase (GCK) from glucokinase regulatory protein (GCKR) (Brown et al., 1997; McGuinness and Cherrington, 2003). While the same mechanism may apply to β-cells, it is less likely given the low expression of GCKR in MIN6-K8 cells (Supplementary Table 1). Instead, the observed increase in G6P due to fructose was likely a result of diminished PFK activity. Some studies have reported that fructose is converted to F6P and G6P, which are then oxidized through the pentose phosphate pathway (Sener and Malaisse, 1988; Giroix et al., 1999). However, our findings indicated that [U-^13^C]-fructose was not converted to M6 F6P or M6 G6P. It is possible that differences in experimental conditions, such as sugar concentrations and the species of animals tested, as well as methodological differences, contributed to this discrepancy.

This study provides the first description of fructose-mediated PK inhibition in β-cells. The effect of F1P on PK may be dependent on concentrations and cell types, because it is also reported that liver PK is activated by F1P in liver homogenates (Eggleston and Woods, 1970). Nevertheless, our results clearly demonstrated the inhibition of PKM2 by fructose in β-cells, according to the significant accumulation of 2PG and 3PG, which was ameliorated by TEPP-46.

Although TEPP-46 counteracts fructose-mediated PK inhibition, the reason why the insulinotropic effect of TEPP-46 requires intracellular fructose metabolism remains unclear. There was no evidence that ATP production from fructose directly stimulated FSIS, even when TEPP-46 was present, as the whole-cell ATP/ADP ratio visualized by Perceval HR was similar under fructose- and glucose-stimulated conditions. However, our results do not preclude the possibility that local ATP production by PK near the K_ATP_ channel regulates FSIS. Recent studies have proposed that membrane-associated PK directly provides ATP to β-cell K_ATP_ channels and controls glucose-induced channel closure (Lewandowski et al., 2020; Foster et al., 2022; Ho et al. 2023; Merrins and Kibbey, 2024), although this proposal remains controversial (Corradi et al., 2023; Satin et al., 2024; Rutter and Sweet, 2024). A similar mechanism may also participate in FSIS.

Furthermore, lipogenesis-related metabolites, such as diacylglycerol, free fatty acid, and fatty acid CoA, are potential metabolic coupling factors (MCFs) in FSIS (Prentki et al., 2013). Fructose-derived metabolites are preferentially used for de novo lipid synthesis (Herman and Samuel, 2016; Silva et al., 2019). As our metabolic analysis did not cover these pathways, more comprehensive metabolic profiling is required to elucidate the link between intracellular fructose metabolism and FSIS.

Excessive fructose consumption has been linked to several health issues, including obesity, metabolic dysfunction-associated steatotic liver disease (MASLD), and type 2 diabetes (Hannou et al., 2018; Taskinen et al., 2019). It has been demonstrated that FSIS is impaired in the islets of NSY.B6-Ay/a, a mouse model of obese diabetes, suggesting that the deleterious effects of fructose on glucose tolerance are counterbalanced by increased insulin secretion. Surprisingly, these diabetic islets became responsive to fructose when treated with TEPP-46. Furthermore, a recent study showed that pyruvate kinase activators improve both insulin sensitivity and insulin secretion in high-fat diet-fed mice in vivo (Abulizi et al. 2020). Given these findings, additional research is necessary to evaluate TEPP-46’s potential as a therapeutic agent for fructose-induced metabolic disorders.

## 8 LIMITATIONS OF THE STUDY

In this study, we utilized fructose concentrations that are in line with recent investigations (Kyriazis et al., 2012; Bartley et al., 2019), but are not confirmed to be physiologically relevant. We recognize that insulin secretory response to ingested fructose is determined by blood fructose levels in vivo. However, there are considerable variations in the reported blood fructose levels, ranging from sub-millimolar values (Sugimoto et al., 2010; Patel et al., 2015) to more than 10 millimolar (Hui et al., 2009), which is likely due to differences in the subjects and quantification methods. Future studies should aim to establish dose-response relationships for fructose-induced insulin secretion in vivo to better understand the physiological implications of our findings.

## 9 AUTHOR CONTRIBUTIONS

Conceptualization, N.M.; Methodology, N.M..; Investigation, N.M. and R.M.; Resources – T.O., and N.Y.; Writing – Original Draft: N.M.; Writing – Review & Editing: N.M., R.M., Y.S., K.S.., Y.M., T.O.., N.Y.., Y.Y., and A.S.; Data Curation: N.M.; Visualization: N.M.; Supervision: K.S., Y.Y., and A.S.; Funding Acquisition: N.M. and A.S.

## 10 ACKNOWLEDGEMENTS

The authors extend their gratitude to President Yutaka Seino of Kansai Electric Power Hospital for his general support in this research. The authors thank Yoshikazu Hoshino of Hoshino Laboratory Animals, Inc. for his assistance with the NSY.B6 mice. The authors thank Shihomi Sakai, Asami Yamaguchi, and Megumi Akiyama for their excellent technical assistance.

## 11 DATA AVAILABILITY

The RNA sequencing data used for Supplementary Table 1 are available from the DDBJ Sequence Read Archive with the accession number DRA006332 (Hashim et al., 2018). Other data supporting the findings of this study are available from the corresponding author upon reasonable request.

## 3 ABBREVIATIONS

2PG: 2-phosphoglycerate
3PG: 3-phosphoglycerate
DHAP: dihydroxyacetone phosphate
FBP: fructose 1,6-bisphosphate
F1P: fructose 1-phosphate
FSIS: fructose-stimulated insulin secretion
G6P: glucose 6-phosphate
F6P: fructose 6-phosphate
GA3P: glyceraldehyde 3-phosphate
GCK: glucokinase
GCKR: glucokinase regulatory protein
GSIS: glucose-stimulated insulin secretion
IP1: inositol 1-phosphate
K_ATP_ channel: ATP-sensitive K^+^ channel
KHK: Ketohexokinase
MASLD: metabolic dysfunction-associated steatotic liver disease
OAA: oxaloacetate
PEP: phosphoenolpyruvate
PK: pyruvate kinase
PLC: phospholipase C

## SUPPLEMENTARY MATERIALS

**Supplementary Figure 1.**
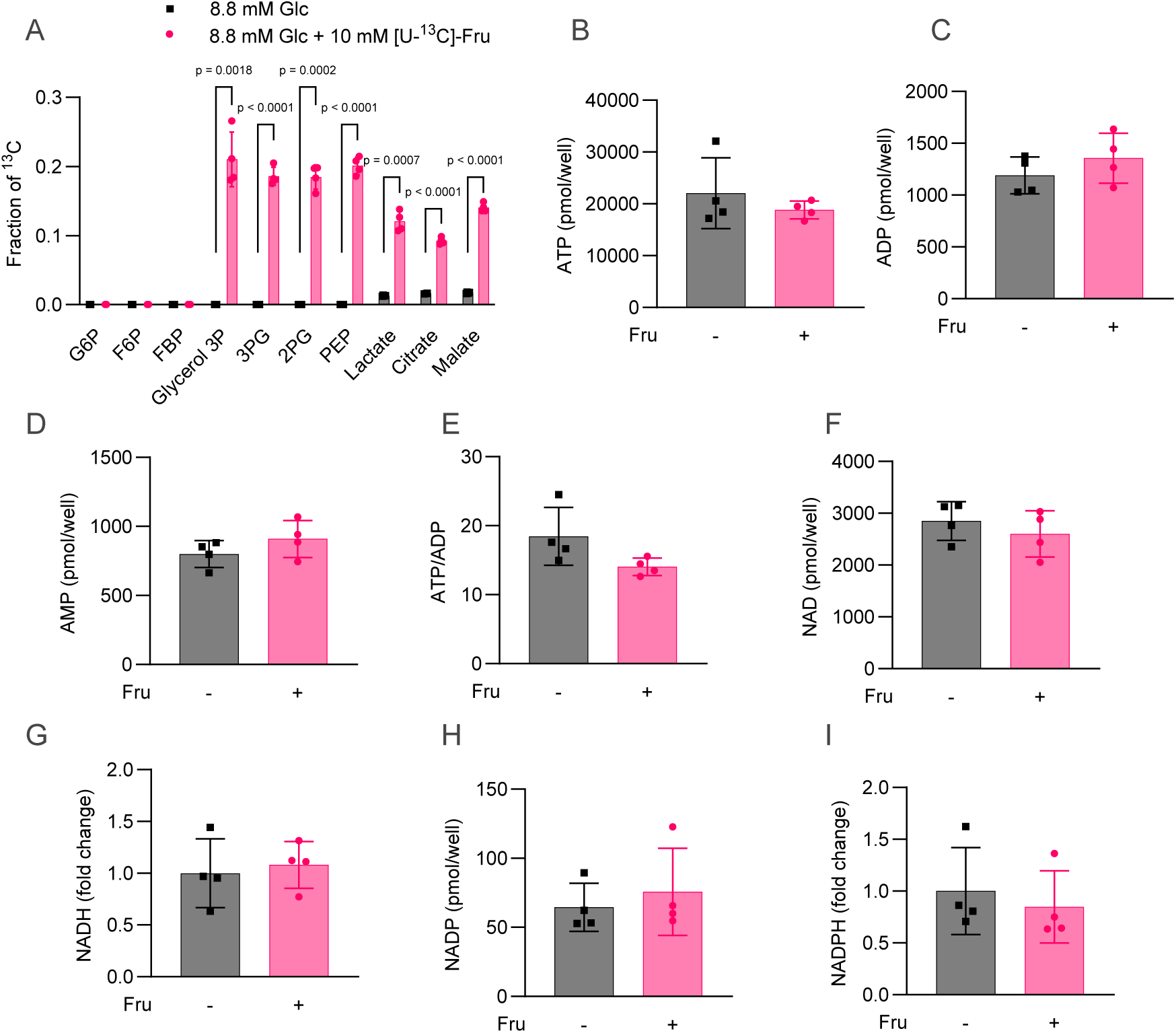
[U-^13^C]-fructose tracing in MIN6-K8 cells. (A) Fraction of ^13^C in each intermediate. (B) –(I) Total metabolite content and their ratio. All experiments were performed using MIN6-K8 cells. n = 4. Data are presented as the mean ± SD. Statistical comparisons were made using Welch’s unpaired two-tailed t-test. See also Figure 2 for the experimental details.

**Supplementary Figure 2.**
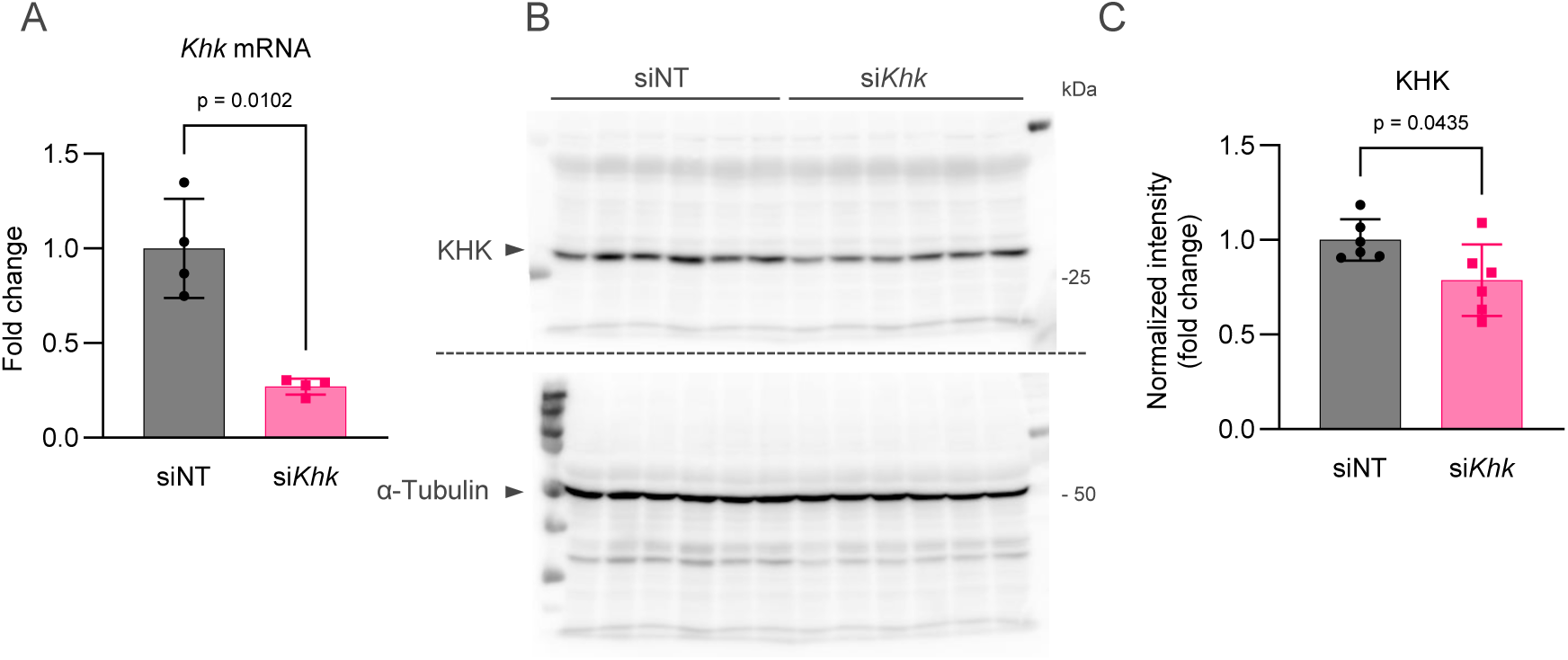
(A) Knockdown efficiency of *Khk* was assessed by RT-qPCR. mRNA levels were normalized to those in siNT (non-targeting siRNA)-treated cells. n = 4. (B) –(C)Knockdown efficiency of KHK protein assessed by immunoblotting. (B) The membrane was first probed for KHK and subsequently re-probed for α-tubulin. The corresponding bands are indicated by black triangles. (C) The average intensity of KHK was normalized to that of α-tubulin and expressed as fold-change. n = 6. All experiments were performed using MIN6-K8 cells. Data are presented as the mean ± SD. Statistical comparisons were made using Welch’s unpaired two-tailed t-test.

**Supplementary Figure 3.**
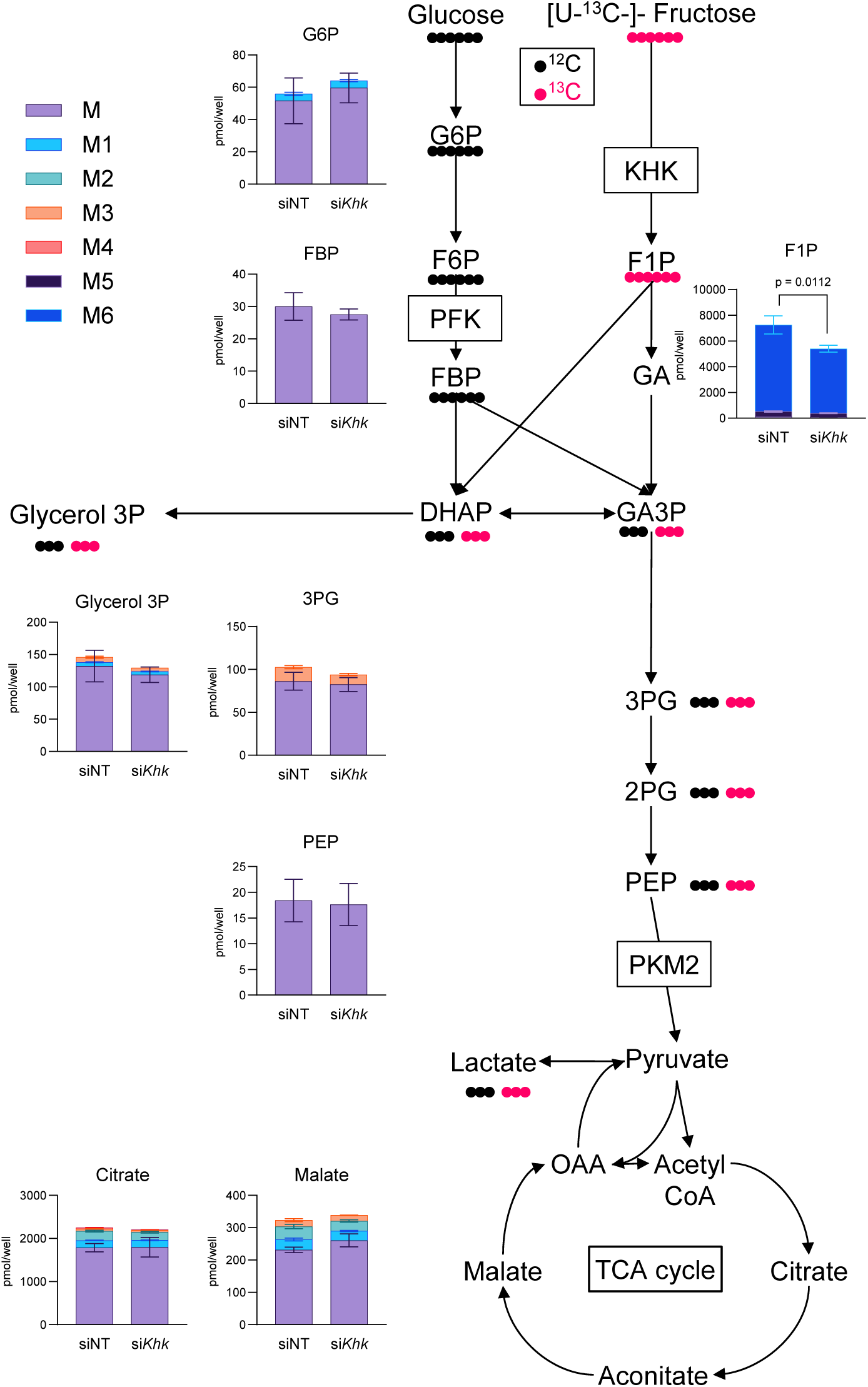
Effect of Khk knockdown on [U-^13^C]-fructose tracing in MIN6-K8 cells. The total content and isotopic distribution of each intermediate are presented as stacked bar graphs along with a schematic overview of the metabolic fates of glucose and [U-^13^C]-fructose. n = 4. Statistical comparisons were made between the total content using Welch’s unpaired two-tailed t-test.

**Supplementary Figure 4.**
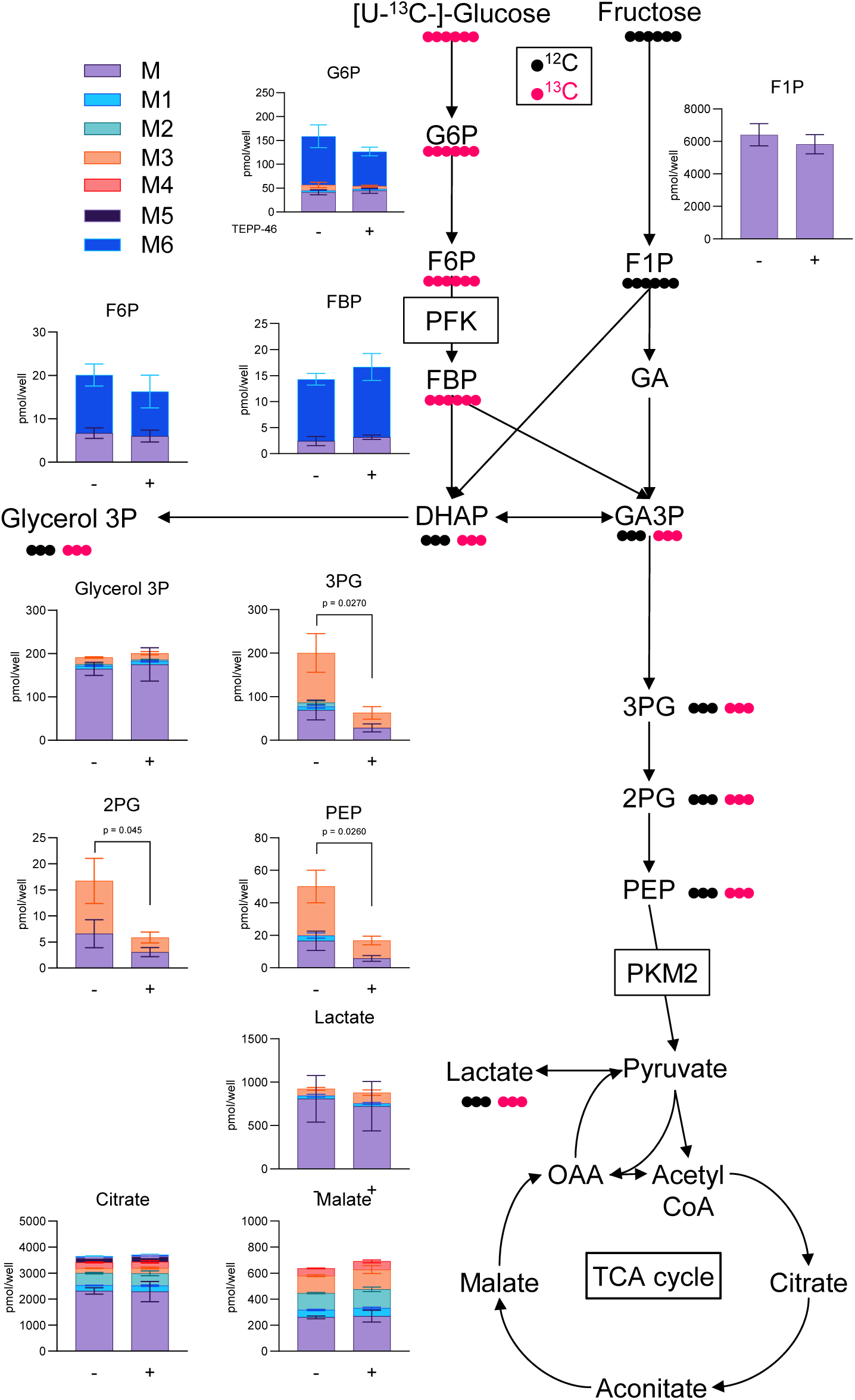
Effect of TEPP-46 on [U-^13^C]-glucose tracing in MIN6-K8 cells. The total content and isotopic distribution of each intermediate are presented as stacked bar graphs, along with a schematic overview of the metabolic fates of [U-^13^C]-glucose and fructose. n = 4. Statistical comparisons were made between the total content using Welch’s unpaired two-tailed t-test.

**Supplementary Figure 5.**
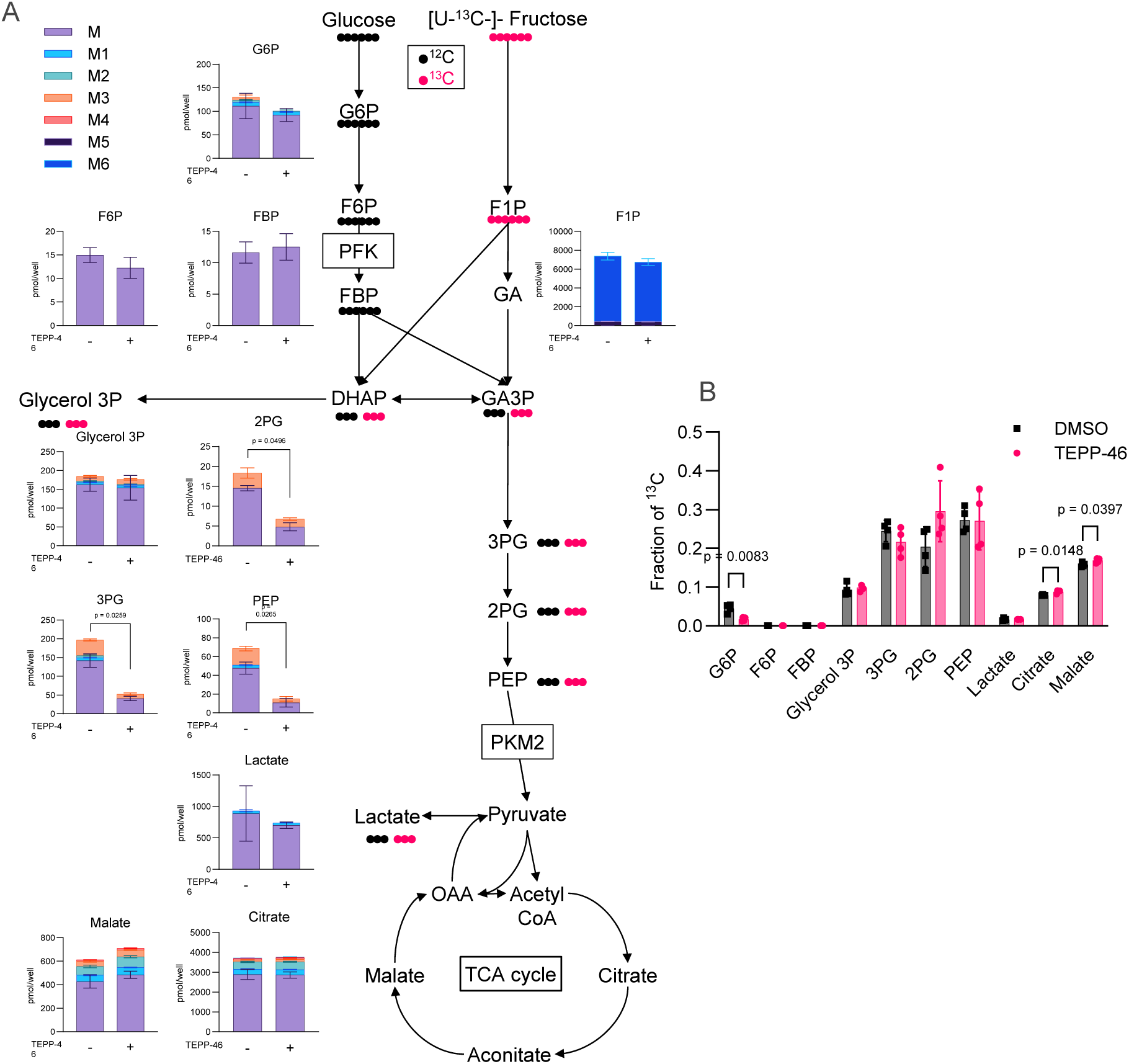
Effect of TEPP-46 on [U-^13^C]-fructose tracing in MIN6-K8 cells. n = 4. (A) The total content and isotopic distribution in each intermediate are presented as stacked bar graphs. Statistical comparisons were made between the total content. (B) Fraction of ^13^C in each intermediate. Statistical comparisons were made using Welch’s unpaired two-tailed t-test.

**Supplementary Table 1.**
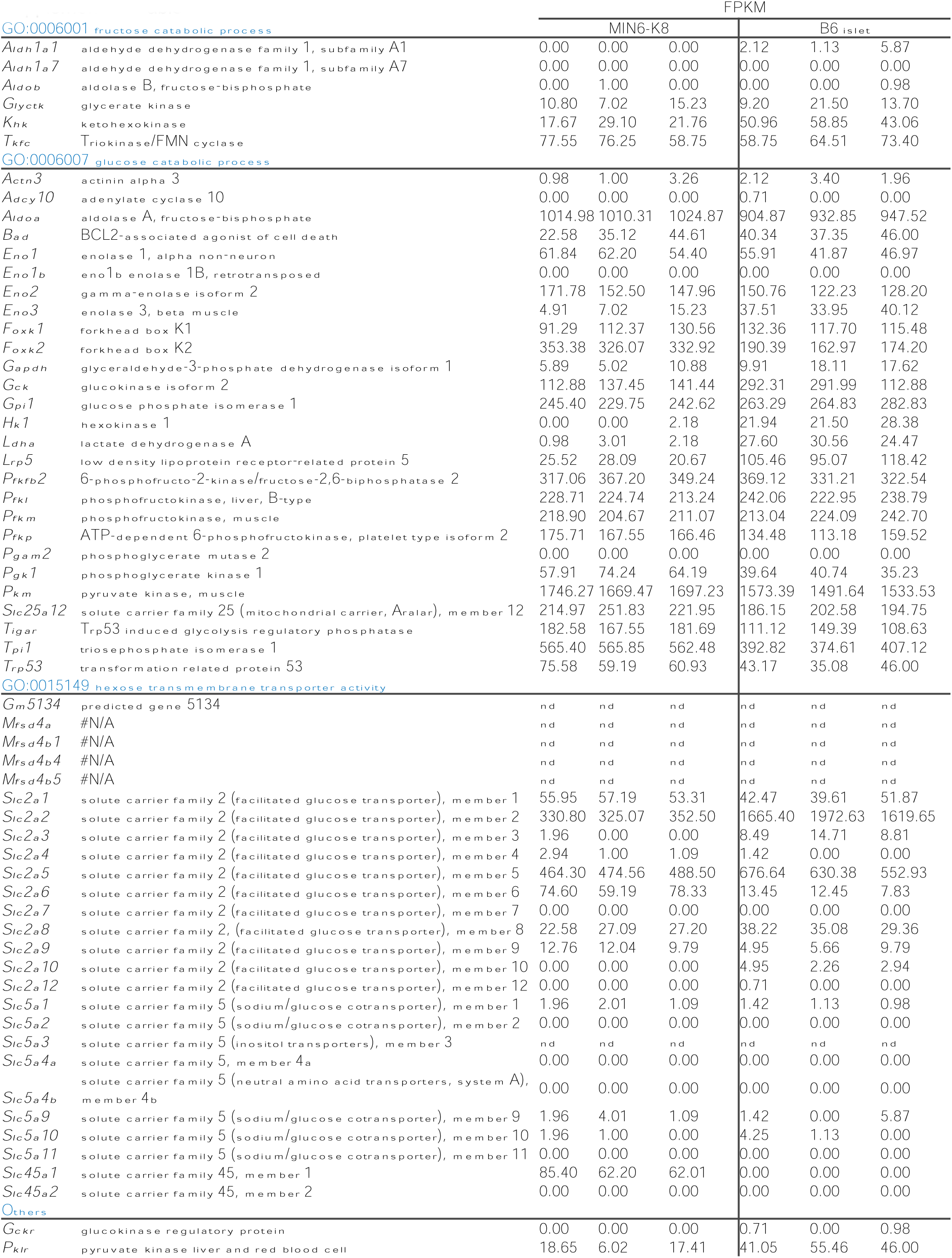
Expression profiles of glucose/fructose catabolism-related genes in MIN6-K8 cells and B6 mouse islets. Source data: DDBJ Sequence Read Archive (DRA) accession number DRA006332 (Hashim et al., 2018). Genes were grouped according to the indicated Gene Ontology (GO) terms. n = 3 for each. nd, not detected.

